# Precise generation of bystander-free mouse models with ABE9-SpRY

**DOI:** 10.1101/2025.10.07.680931

**Authors:** Jun Kai Ong, Sayari Bhunia, Beate Hilbert, Vanessa Kirschner, Sascha Duglosz, Frank Zimmermann, Marc Freichel, Alex Cornean

**Affiliations:** Institute of Pharmacology, Heidelberg University, Heidelberg, 69120, Germany; DZHK (German Center for Cardiovascular Research), Partner Site Heidelberg/Mannheim, Heidelberg University, Heidelberg, 69120, Germany; Heidelberg Biosciences International Graduate School (HBIGS), Heidelberg, 69120, Germany; Interfacultary Biomedical Faculty (IBF), Heidelberg University, Heidelberg, 69120, Germany; Medizinische Klinik II, Uniklinikum Würzburg, Würzburg, 97078, Germany

**Author notes:** To whom correspondence should be addressed. Correspondence may also be addressed to.

**Keywords:** adenine base editing, precision editing, mouse modelling, CRISPR/Cas9, hiPS cells

## Abstract

Point mutations cause many genetic disorders, but modelling them in organisms is technically challenging. Creating mouse models that mimic these mutations is crucial for establishing a causal relationship between mutations and disease phenotype, thereby supporting the development of therapeutic strategies. Adenine base editors (ABEs) can correct single-nucleotide variants (SNVs) in disease modelling without double-stranded breaks (DSBs) or donor DNA, achieving higher product purity than traditional Cas9 methods. Earlier ABE techniques faced issues like limited targetability, bystander editing, and off-target effects. By combining two editor advancements, we introduced and tested ABE9-SpRY, an improved ABE variant fused with a PAM-flexible SpRY-Cas9 nickase. Our results show that ABE9-SpRY effectively generates three out of four targeted A-to-G mutations in mouse embryos, with significantly fewer off-target effects than ABE8e-SpRY, achieving desired editing efficiencies of up to 96% in individual adult founder mice.ABE9-SpRY also enhances product purity in mouse embryos and human induced pluripotent stem cells (hiPSCs) compared to ABE8e-SpRY. Our findings showcase ABE9-SpRY’s precision and versatility, highlighting it as a powerful tool for accurate *in vivo* point mutation modelling.

## INTRODUCTION

Disease models have been indispensable in advancing biomedical research by enabling the study of complex physiological and biological interactions that are difficult to replicate *in vitro* ^1^. The mouse has emerged as the dominating biomedical model due to its genetic, biological, and behavioural similarities to humans, along with the availability of advanced genetic tools for manipulation ^2^. Moreover, mouse models with disease-relevant mutations or mutations in a given drug receptor have tremendous value for the development and validation of the mode of action of new therapies ^3-5^. In parallel, hiPSCs have also become a powerful *in vitro* model, providing a human-specific system for studying disease mechanisms, drug responses, and genetic modifications in a physiologically relevant context ^6^.

The recent development of precise CRISPR/Cas-based gene editing tools has significantly expanded the possibilities for creating genetically engineered mouse models. Notably, the CRISPR/Cas9 system has enabled precise gene modifications by inducing DSBs at specific genomic loci ^7^. Its efficiency, widespread adaptation and simplicity have made CRISPR/Cas9 the most widely used system for gene editing. However, the CRISPR/Cas9 system has its limitations. The requirement for DSBs introduces risks such as off-target effects and the formation of insertions or deletions (indels) due to the predominant non-homologous end-joining (NHEJ) repair pathway, which can unintentionally disrupt the targeted site or introduce unintended mutations elsewhere ^8,9^. Likewise, larger deletions ^10^ and chromosomal aberrations ^11^ have been reported to occur at significantly higher frequencies with DSB-inducing genome editing approaches.

To address the limitations of DSBs, DNA base editing has emerged, which is catalysed by a new class of tools known as base editors ^12,13^. Base editors allow single nucleotide conversions without needing DSBs or donor DNA templates. ABEs are of particular interest for introducing A-to-G (or T-to-C) base transitions ^12^, which are relevant to a wide range of genetic disorders. For example, G•C to A•T mutations account for approximately 47% of the most common pathogenic SNPs in the ClinVar database ^14^. By fusing a Cas9 nickase (Cas9n) with a deaminase enzyme that converts adenine to inosine, then read as guanine by the cell, ABEs have been used to install point mutations with high efficiency and precision ^12,15^.

ABEs, therefore, hold immense promise for enabling the rapid and precise generation of disease models in animal and cell models, especially in mice. However, current ABE approaches in mice are limited by bystander editing, issues of targetability and off-target (OT) editing **(Fig. 1**). Bystander editing occurs when nucleotides within the editing window of the deaminase, other than the target site(s), are also edited ^14^, leading to unwanted mutations and making it difficult to study the disease mutations specifically. Canonical ABEs are also limited by their dependency on the NGG protospacer adjacent motif (PAM) recognised by the commonly used *Streptococcus pyogenes* Cas9 nickase, restricting the range of targetable sites in the genome ^12^. Both Cas-dependent, guided by the sgRNAs’ base pairing properties to similar genomic regions ^16,17^, and Cas9-independent off-target editing, based on deaminase activity on adenines in single-stranded DNA regions outside the targeted site ^18,19^, have raised serious concerns. These limitations have driven efforts to improve ABEs through directed evolution and rational design. Next-generation variants, such as ABE8e ^20^, exhibit enhanced deaminase activity, whereas ABE9 has increased specificity due to a narrower editing window ^21^. Additionally, engineered Cas9 variants like SpRY, which have relaxed PAM requirements, have expanded the range of targetable sequences, allowing for more flexible editing across the genome ^22^. We therefore aimed to combine the specificity of the ABE9 deaminase with the flexibility of SpRY, thereby overcoming all three challenges and benchmarking these against ABE8e-SpRY ^23^, to establish four different mouse models with precise point mutations in the ion channels TPC1, TPC2, and TRPM4. These channels play critical roles in physiological processes, and mutations in these channels have been linked to diseases, particularly affecting hepatic and cardiovascular tissues ^24-28^.

**Figure 1.**
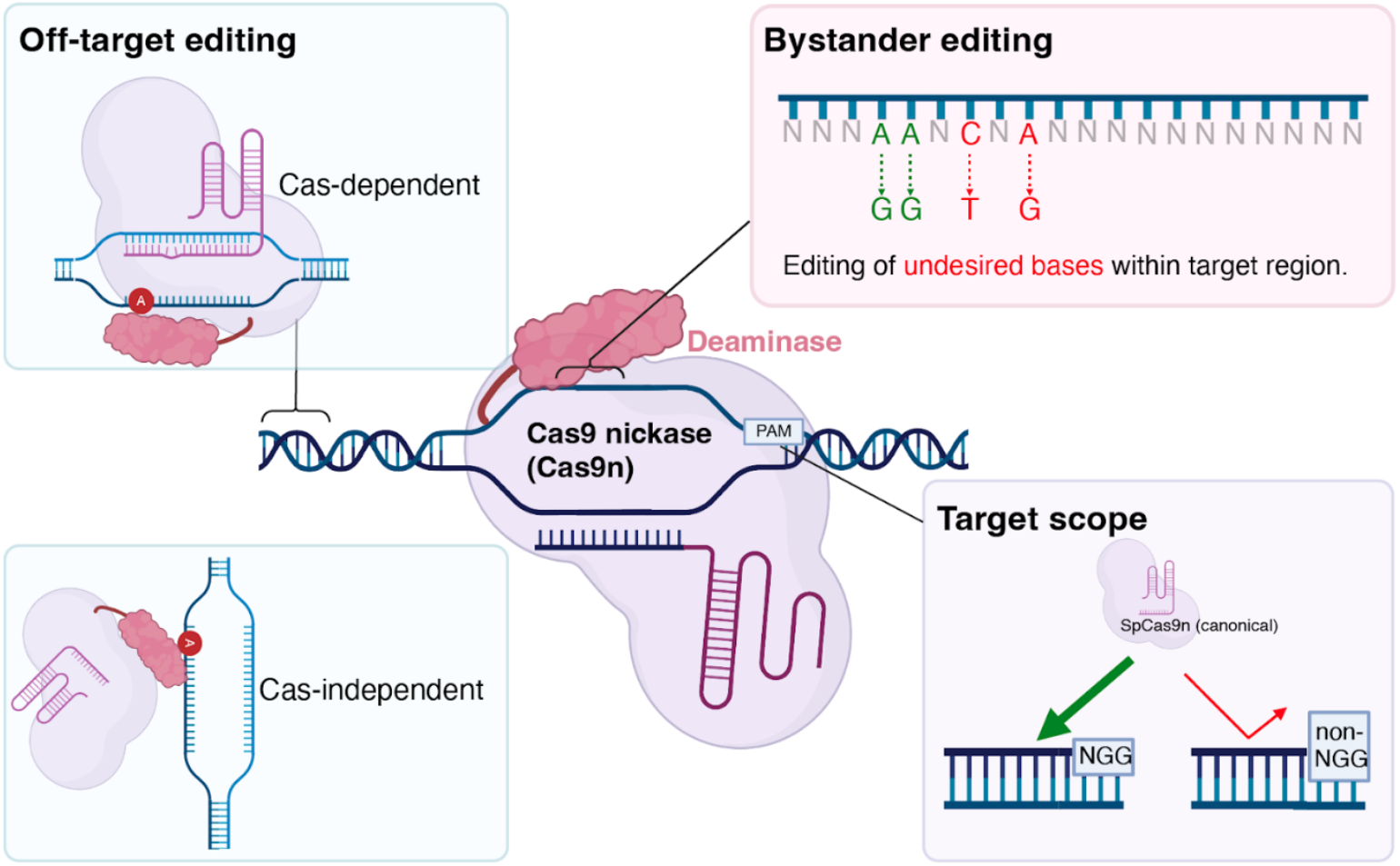
Limitations of disease modelling with ABEs. Schematic of limitations of ABEs in generating disease models. ABEs can have Cas-dependent or -independent off-target editing. Besides targeted adenines, ABEs also have bystander activity on non-target bases. SpCas9 employed in ABEs recognises a 3’ NGG PAM while does not bind optimally, or at all, to non-NGG PAM sites. This Figure was partially prepared with BioRender.

To streamline the generation of disease models in mice and substantially reduce the number of required animals, we established a workflow to characterise and employ ABE9-SpRY for the rapid introduction of desired mutations. After initial benchmarking and strategy design, we cloned and tested the desired sgRNA/ABE-SpRY combinations in N2a cells, followed by pooled injection of synthetic sgRNAs and SpRY-ABEs mRNA into mouse zygotes and analysis of all target and selected OT sites in E14 embryos. Our strategy enabled generating two point mutation mouse models in the absence of bystander editing and with high efficiency for *Tpc1*^*I486T*^ and *Trpm4*^*L903P*^, which can, due to its broad targetability, be used to create a substantial fraction of A-to-G-based mouse models.

## RESULTS

### Efficient and precise A-to-G base editing with ABE9 in human cells

The most widely used ABE variant today is ABE8e, recognised for its high deaminase activity and processivity ^20^. However, the increased catalytic activity coincides with a significant broadening of the editing window, expanding from adenine positions A4-A8 in ABEmax to A3-A10 in ABE8e ^29^, as well as higher levels of off-target DNA and RNA editing ^20,21,29^. Therefore, this rise in unwanted on-target bystander and off-target editing raises concerns regarding the safety of ABE8e for potential clinical applications ^29^. To address these safety concerns, several approaches have frequently been employed in the past (**Table 1**): introducing structure-guided or evolved mutations in the deaminase domain ^15,20,21,29-33^, splitting the editor ^34,35^, using low mismatch tolerance Cas variants ^36^, and fusing the ABE with an effector domain ^37^.

**Table 1.**
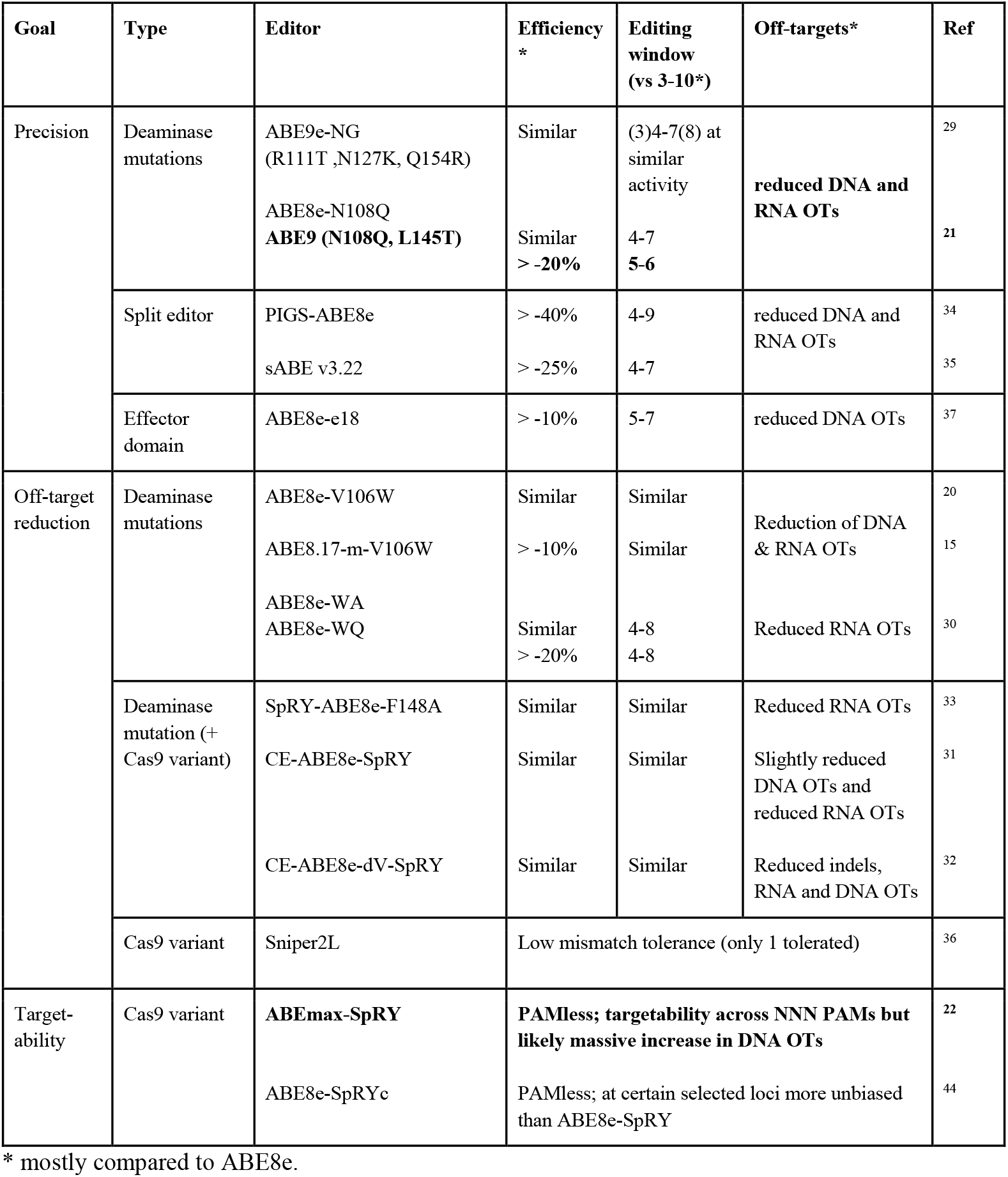
Overview of available approaches to overcome precision, off-target (OT) and targetability issues.

Among all these options, ABE9, which was derived from ABE8e by introducing two point mutations N108Q and L145T in the deaminase domain, appeared to be the most suitable due to its narrowest editing window to date, located between protospacer positions 5-6, and its strong properties for reducing off-target effects in DNA and RNA ^21^. To this end, we compared ABE8e and ABE9 at four previously reported ABE target sites in HEK293T cells to confirm their properties (**Supplementary Fig. 1**). Indeed, the editing window of ABE9 was specific to positions A5 and A6, with some minimal activity at A4, whereas ABE8e was highly active between A3 and A8 with an extended activity shoulder to A13. Overall, the average activity at A5 was reduced from 43.6 ± 5.9% (mean±s.d.%) in ABE8e to 34.5 ± 8.8% in ABE9, corresponding to an average reduction of 21% in editing activity, in line with previous work ^21^ (**Supplementary Fig. 1**). We were, therefore, able to confirm the potential of ABE9 and, encouraged by this, proceeded to apply its precision to our loci of interest.

### ABE9-SpRY is a precise and versatile editor in mouse cells

We first designed the sgRNAs for the four loci to position the target adenines within the optimal editing windows of the deaminases of ABE8e and ABE9, respectively (**Fig. 2a**). However, the canonical architectures of PAM-interacting motifs in both ABEs, when fused to the standard *Sp*Cas9n, proved unsuitable due to the lack of the NGG PAM sequence.

**Figure 2.**
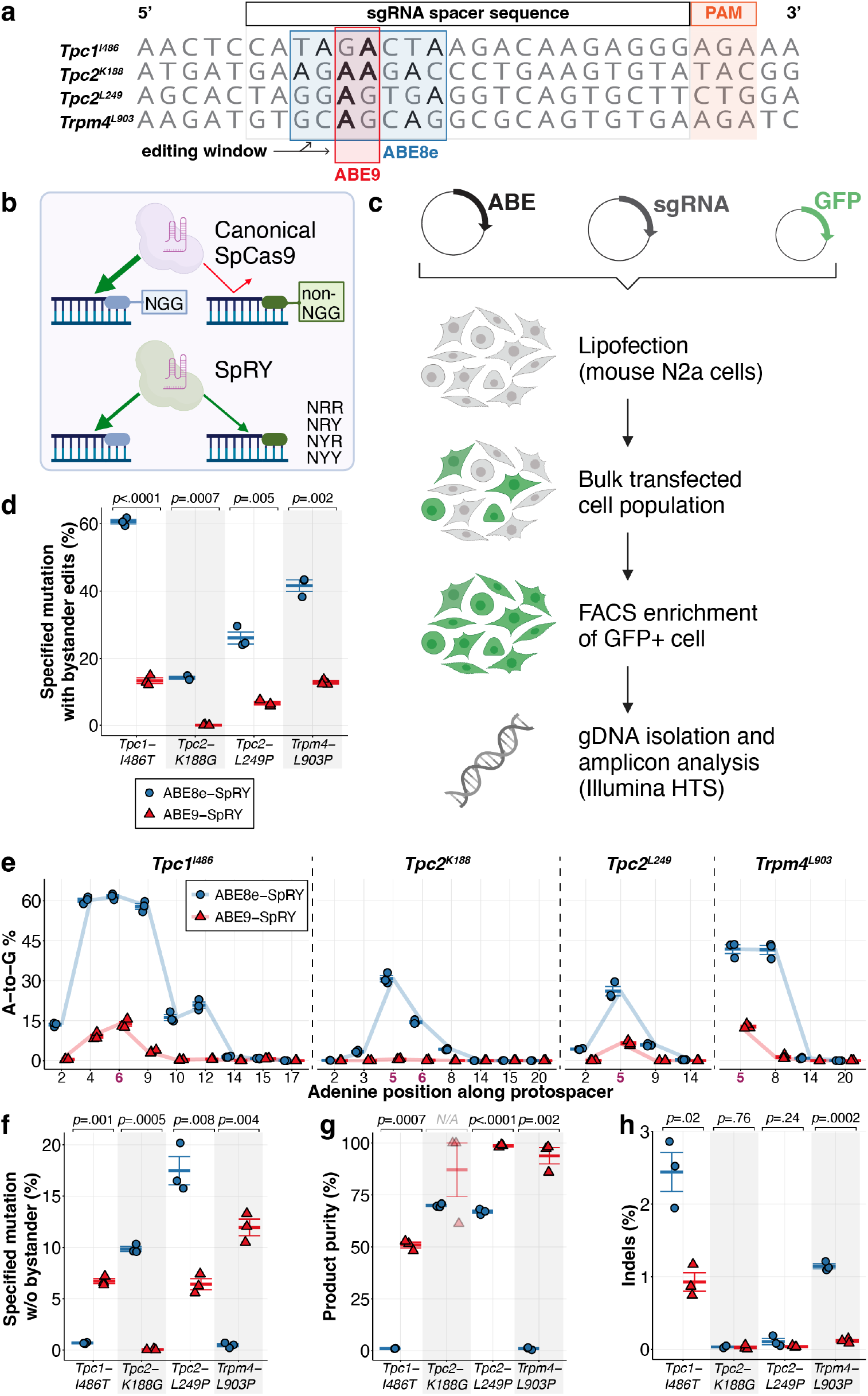
Evaluation of ABE-SpRY constructs in target sites of interest in mouse N2a cells. (**a**) Design of sgRNAs for optimal adenine base editing. Target adenines (bolded in red) for each locus are positioned within the optimal editing windows of ABE8e and ABE9 deaminases with the corresponding PAM sites shown. (**b**) The Cas9 variant SpRY can target non-NGG PAM sites, unlike the wild-type SpCas9. (**c**) N2a cells were co-transfected with three plasmids: the ABE plasmid, sgRNA plasmid, and GFP plasmid. GFP-positive cells were sorted, and genomic DNA (gDNA) was extracted for subsequent targeted sequencing analysis. (**d**) Maximum editing efficiency observed for the target adenine using ABE8e- or ABE9-SpRY, regardless of bystander editing. (**e**) A-to-G editing efficiencies of ABE8e and ABE9-SpRY at each adenine position along the protospacer region for the four target sites. The positions of adenines of interest are bolded in red. (**f**) Editing efficiency of the target adenine using ABE8e- or ABE9-SpRY, considering only reads without bystander editing or indels. (**g**) Product purity, calculated as the proportion of correctly edited sequences (F) relative to the total editing events (D), expressed as a percentage (F/D × 100%). (**h**) Frequency of indels at each of the four target sites. Data is presented as mean ± s.e.m., and individual data points are shown for three biological replicates. Parts of the schematics in (**c**) were created with BioRender.com.

To address this limitation, we next considered alternative Cas9 variants that have different or relaxed PAM requirements, including *Sa*Cas9, xCas9, *Sp*Cas9-NG, *Sp*Cas9-VQR, and SpRY ^22,38-43^. Instead of an “on-demand” context-dependent approach, we developed a versatile editing strategy that can be applied to all four loci of interest by utilising a single construct and the appropriate sgRNA. SpRY ^22^ and SpRYc ^44^ have emerged as potentially universal Cas9 variants with PAMless targetability across NNN PAMs (**Table 1**). SpRY stands out for having the broadest and most relaxed PAM compatibility and has previously been tested with ABE8e variants ^31-33^. We fused a SpRY nickase (in the following simply “SpRY”, bears these substitutions compared to *Sp*Cas9 nickase: A61R/L1111R/D1135L/S1136W/G1218K/E1219Q/N1317R/A1322R/R1333P/R1335Q/T1337R) to the TadA domains of ABE8e or ABE9 to streamline the experimental workflow and maximise the potential for successful genome editing outcomes (**Fig. 2b**). These SpRY base editors can, in theory, effectively target spacers with NRN (G or A) PAMs, and to a lesser degree, NYN (T or C) PAMs ^22^.

Since the compatibility of deaminase domains with Cas9 variants is not universal, and their efficiency and editing activity can vary based on the specific Cas9 variants used ^15,45,46^, an initial evaluation was necessary. Due to the lack of published reports on ABE9-SpRY, we compared it to ABE8e-SpRY using a plasmid-based lipofection approach alongside the respective sgRNAs for the four loci of interest in mouse Neuro-2a (N2a) cells (**Fig. 2c**).

Deep sequencing of transfected cells revealed that the desired editing occurred at all four targets for ABE8e-SpRY, ranging from 14.1 ± 0.7% to 60.6 ± 1.2%. In contrast, only three targets showed targeted A-to-G editing with ABE9-SpRY, ranging from 6.5 ± 0.9% to 13.2 ± 1.5% across all tested sites (**Fig. 2d**). While ABE8e-SpRY displayed modest editing of 14.1 ± 0.7% at the *Tpc2-K188* locus, no desired editing was noted with ABE9-SpRY. A closer examination of editing at adenines within the sgRNA target sites confirmed the previously observed broad activity window of ABE8e-SpRY, which included 13.5 ± 0.5% editing at A2 and 20.8 ± 0.6% editing at A12 at the *Tpc1-I486* locus (**Fig. 2e**). The highest ABE8e-SpRY activities were observed at A5 (*Tpc2-K188, Tpc2-L249, Trpm4-L903*) and A6 (*Tpc1-I486)*, with 30.8 ± 0.5%, 26.1 ± 6.5%, 41.9 ± 12.8%, and 61.6 ± 12.7% A-to-G editing, respectively. By contrast, ABE9-SpRY demonstrated the enhanced precision previously observed for ABE9, with the activity window spanning from A4 to A9 at the *Tpc1-I486* locus, showing 9.1 ± 1.3% and 3.3 ± 0.5% editing, respectively. We also measured the highest efficiencies for ABE9-SpRY at A5 and A6, respectively, with 6.5 ± 0.9%, 12.8 ± 0.8%, and 13.7 ± 1.7% efficiency for *Tpc2-L249, Trpm4-L903*, and *Tpc1-I486*, respectively. This narrower editing window of ABE9-SpRY also led to a larger fraction of edited alleles containing no bystander mutations compared to ABE8e-SpRY (**Fig. 2f**), as indicated by the substantially higher product purity, ranging from 50.9 ± 2.4% to 98.6 ± 0.6%, compared to 1.1 ± 0.1% to 69 ± 0.8% for ABE8e-SpRY (**Fig. 2g**). Notably, the desired *Tpc1*^*I486T*^ and *Trpm4*^*L903P*^ mutations could not be introduced without bystanders using ABE8e-SpRY (0.7 ± 0.1% and 0.5 ± 0.3%), whereas ABE9-SpRY achieved 6.7 ± 0.4% and 11.9 ± 1.4% precise editing at these sites, respectively. Lastly, ABE9-SpRY demonstrated reduced indel frequencies compared to ABE8e-SpRY across all four sites, peaking at 0.9 ± 0.2% (*Tpc1-I486*) (**Fig. 2h**).

Taken together, these *in vitro* results for ABE target sites in mouse cell line indicated that ABE9-SpRY can precisely edit three out of four desired target sites, two of which could not be generated with ABE8e-SpRY due to bystander mutations. Therefore, we next decided to apply these approaches *in vivo*, testing editing efficiencies in mouse embryos.

### Multiplexed, precise ABE9-SpRY mouse embryo editing

Adenine base editing in mouse zygotes has been successfully performed using a combination of ABE mRNA and either *in vitro* transcribed or chemically synthesised modified sgRNAs, injected into the cytoplasm or pronucleus of the zygote ^37,47-52^. Indeed, these efforts have been highly efficient in achieving nearly 100% editing using ABE7.10 ^47-49^. However, precision issues were apparent, as only 1.3% to 20% of desired edits occurred within an overall A-to-G efficiency of 95% in the protospacer^48^.

To evaluate the *in vivo* editing efficiency of our approach in mouse embryos, we microinjected a pool of four synthetic sgRNAs along with either *in vitro* transcribed ABE8e-SpRY or ABE9-SpRY mRNA into the cytoplasm of mouse zygotes. The injected zygotes were transferred to pseudopregnant surrogate mothers, isolated on embryonic day 14, and their genomic DNA was extracted for targeted deep sequencing analysis (**Fig. 3a**).

**Figure 3.**
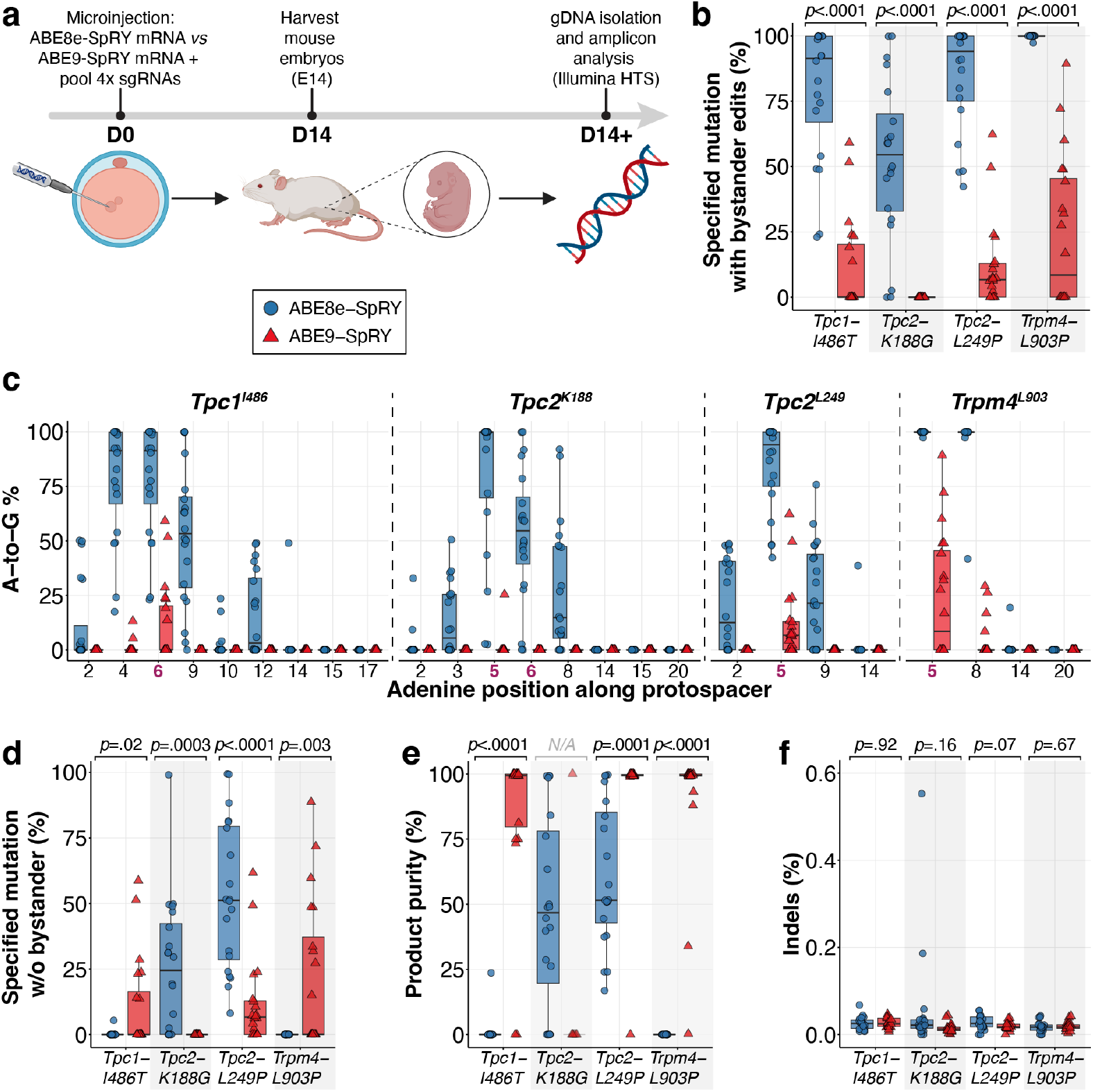
Evaluation of ABE-SpRY constructs in target sites of interest in mouse embryos. (**a**) Schematic of generation of genetically edited mouse embryos. (**b**) Maximum editing efficiency observed for the target adenine, regardless of bystander editing. (**c**) A-to-G editing frequencies in twenty ABE8e or ABE9-SpRY edited mouse embryos at each adenine position along the protospacer region for the four target sites. Target adenines are bolded in red. (**d**) Maximum desired editing efficiency excluding cases with bystander editing. (**e**) Product purity, calculated as the proportion of correctly edited sequences (D) relative to the total editing events (B), expressed as a percentage (D/B × 100%). (**f**) Frequency of indel occurrence at the four target sites. Data are displayed as standard boxplots following the Tukey convention, including the median (horizontal line), and individual data points are shown for 20 independent biological replicates. Parts of the schematics in (**a**) were created with BioRender.com.

Our analysis indicated that ABE8e-SpRY effectively edited target adenines at all four sites, achieving editing efficiencies of up to 100%. The average editing efficiencies were 78.9 ± 25.7% at *Tpc1-I486* [median (IQR), 91.4% (67.0-99.9%)], 52.6 ± 30.8% at *Tpc2-K188* [median (IQR), 54.5% (32.9-70.1%)], 84.5 ± 20.2% at *Tpc2-L249* [median (IQR), 94.1% (75.1-99.9%)], and 99.8 ± 0.6% at *Trpm4-L903* [median (IQR), 99.9% (41.7-100.0%)], (**Fig. 3b**). Editing by ABE9-SpRY was observed at *Tpc1-I486T, Tpc2-L249*, and *Trpm4-L903P*, achieving efficiencies of up to 59.1%, 62.3%, and 89.3%, with corresponding averages of 11.0 ± 18.1% [median (IQR), 0.04% (0.02-20.26%)], 11.9 ± 16.8% [median (IQR), 6.71% (0.04-12.95%)], and 23.7 ± 28.6% [median (IQR), 8.50% (0.03-48%)], respectively.

A closer examination of adenine base editing across the protospacer sequences revealed a pattern consistent with previous *in vitro* findings. Despite the high editing efficiency of ABE8e-SpRY, its activity was broadly distributed, targeting adenines from positions A2 to A14 across all tested loci (**Fig. 3c**). In contrast, ABE9-SpRY exhibited a narrower editing window, concentrating edits primarily within positions A4 to A8. Notably, low-level editing was detected in a single embryo at the *Tpc2-K188* locus, where ABE9-SpRY had shown no *in vitro* activity. This editing occurred at only one of the two targeted adenines, resulting in a 25.2% conversion that introduced a p.K188E substitution instead of the intended p.K188G mutation.

ABE9-SpRY exhibited a higher rate of desired edits with no bystander edits at the *Tpc1-I486* locus (10.7 ± 17.5% vs 0.3 ± 1.2%) and at *Trpm4-L903* (21.3 ± 28.4% vs 0.0 ± 0.0%). In contrast, ABE8e-SpRY showed more specific activity at *Tpc2-K188* (25.5 ± 25.9% vs 0.0%) and *Tpc2-L249* (52.8 ± 27.5% vs 11.8 ± 16.2%) (**Fig. 3d**). Analysing product purity as a measure of precision revealed that ABE9-SpRY generated significantly purer editing outcomes across all active loci: *Tpc1-I486* (85.1 ± 30.6% vs 1.2 ± 5.3%), *Tpc2-L249* (94.6 ± 22.3% vs 60.8 ± 27.1%), and *Trpm4-L903* (90.5 ± 20.8% vs 0.0%) (**Fig. 3e**). Furthermore, ABE9-SpRY produced only low levels of indels, similar to ABE8e-SpRY (**Fig. 3f**).

Only one C-to-T conversion was noted at the *Tpc1-I486* locus in an embryo edited with ABE8e-SpRY, occurring at a frequency of 12.6% at position C6, alongside a single instance of 34.7% unwanted A-to-C editing (**Supplementary Fig. 2a**). Additionally, an unexpected off-target modification was found upstream of the sgRNA binding region at position A-2 of the *Tpc2-L249* locus (**Supplementary Fig. 2b**). This edit appeared in one ABE9-SpRY and one ABE8e-SpRY injected embryo at frequencies of 12.3% and 21.9%, respectively. A possible explanation is the dynamic nature of the Cas9-sgRNA-DNA complex and conformational changes during editing, which may briefly expose adjacent bases to the deaminase domain, leading to unintended deamination events.

These findings suggest that ABE9-SpRY provides enhanced editing purity, though it does so at the expense of locus-dependent activity. To further assess precision beyond these target loci, we next analysed both Cas9-dependent and Cas9-independent off-target editing to more thoroughly characterise the safety profiles of these two editors.

### ABE9-SpRY substantially decreases off-target effects compared to ABE8e-SpRY

Cas-dependent off-target events arise from the tolerance of mismatches between sgRNA and the targeting sequence. Although SpRY can target a broader range of genomic sites, its relaxed PAM tolerance also implies that the potential Cas-dependent off-target sites are substantially expanded, as sgRNA-DNA mismatches become the sole determinant of off-target binding without PAM constraints. Consequently, we identified the top five predicted off-target sites for each locus using ACEofBASEs ^53^ and analysed these sites across three embryos per group.

When analysing the cumulative percentage of reads containing edits in the off-target spacer sequences, we found that the average cumulative A-to-G editing in the off-target protospacer at the *Tpc1-I486* off-target sites varied from 4.1 ± 5.7% to 264.3 ± 15.6% for ABE8e-SpRY, compared to baseline control levels for ABE9-SpRY (**Fig. 4a, Supplementary Fig. 3**). Off-target editing at the highest edited adenines ranged from 4.1 ± 7.0% at OT2 to 27.8 ± 17.4% at OT3 for ABE8e-SpRY (**Supplementary Fig. 3**). Overall, across 20 off-target sites with various PAM sequences, ABE8e-SpRY induced A-to-G edits above control levels at 14 sites, especially at protospacer positions A2 to A20 (**Fig. 4a–d, Supplementary Fig. 3–6**). In contrast, ABE9-SpRY edited just two off-target sites associated with the *Tpc2-L249* locus, reaching 40.0% and 48.1% editing, respectively (**Fig. 4c, Supplementary Fig. 5**). Strikingly, for these two off-target sites, the corresponding embryos’ on-target editing efficiencies at the locus were relatively low, at 7.4% and 10.7%. While off-target effects are known to occur with base editors, this drastic skew towards off-target editing (with a 3-base mismatch) is remarkable. Since these events were observed in just one replicate of ABE9-SpRY, the data indicate that ABE9-SpRY is a notably improved precision tool, though rare, sequence-dependent off-target activity may still occur.

**Figure 4.**
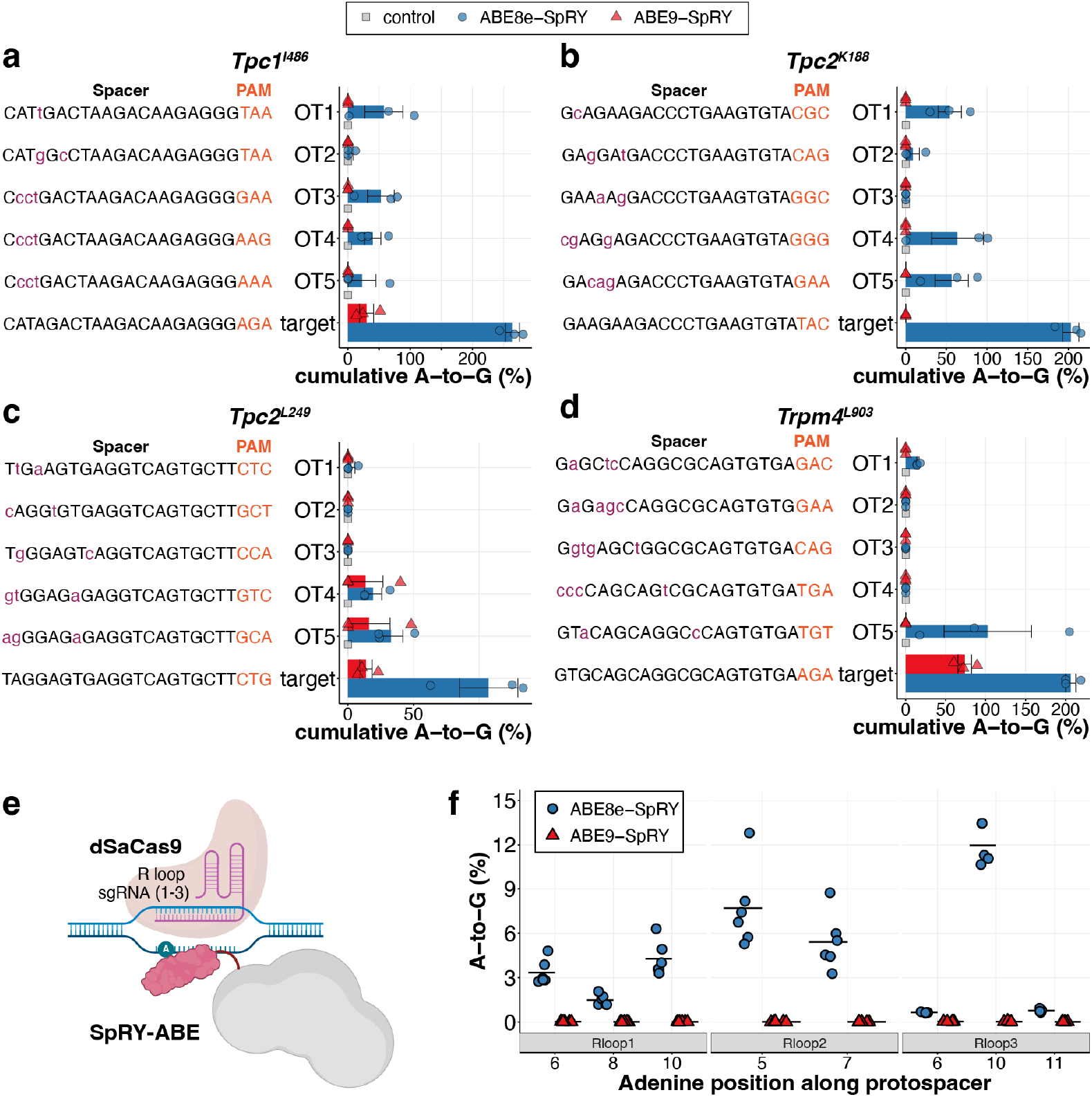
Off-target mutation analysis of ABE-SpRY constructs. (**a-d**) Evaluation of cumulative on- and off-target DNA editing by ABE8e- and ABE9-SpRY in mouse embryos in a Cas9-dependent manner. Data represents mean ± s.e.m. from three biological replicates. (**e**) Diagram of the R-loop assay illustrating how the base editor can target transient non-target R-loops formed by dSaCas9. (**f**) Cas9-independent assessment of off-target DNA editing by ABE8e-SpRY and ABE9-SpRY, using the orthogonal R-loop assay at specific R-loop sites with five or six biological replicates. The black horizontal bar shows the mean at each adenine position grouped by editor. Parts of the schematics in (**e**) were created with BioRender.com.

In addition to Cas-dependent off-target effects, base editors have also been reported to exhibit Cas-independent off-target activity, which is evoked by the deaminase activity of the editors ^18,19^. We therefore asked whether TadA-8e and TadA-9 deaminases cause Cas-independent off-target effects to similar extents, regardless of the Cas9 nickase variant to which they are attached. To this end, we utilised the orthogonal R-loop assay ^54^ (**Fig. 4e**) in HEK293T cells and, after FACS-enrichment of GFP co-transfected cells, observed no editing across eight adenines and three R-loop spacers with ABE9-SpRY, similar to the results reported earlier for ABE9 ^21^. In contrast, ABE8e-SpRY caused substantial editing from 4.3 ± 1.1% (R loop 1), 7.7 ± 2.7% (R loop 2), to 12.0 ± 1.4% (R loop 3). These results highlight the high specificity of ABE9-SpRY despite the increased non-specific binding of SpRY.

### ABE9-SpRY is effective at generating precise *Tpc1*^*I486T*^ and *Trpm4*^*L903P*^ founder mice

Accurate base conversion is crucial for modelling SNVs; therefore, minimising bystander and off-target mutations is essential. Given the negligible levels of bystander editing, low Cas9-dependent off-target editing, and no evidence above background of Cas9-independent off-target editing produced by ABE9-SpRY, we decided to use this editor for generating two individual founder mouse lines harbouring the *Tpc1*^*I486T*^ and *Trpm4*^*L903P*^ mutations. To achieve this, ABE9-SpRY and the locus-specific sgRNAs were injected separately into mouse zygotes. Analysis of adult mice biopsies from 48 putative F0 founder mice revealed robust editing efficiencies (**Fig. 5a,b**), with the precise, bystander, and indel-free p.I486T achieved at 25.6 ± 21.0% efficiency [median (IQR), 26.1% (4.9-44.7%)], and up to 81.4% (**Fig. 5b**). The very low rate of bystander editing at A4 (**Fig. 5b**) resulted in a considerable product purity of 96.4 ± 13.8% [median (IQR), 98.9% (98.8-99.0%)] (**Fig. 5c**). While one individual mouse exhibited an indel frequency of 11.2% at the *Tpc1-I486* locus, overall indel levels were generally minimal (0.3 ± 1.6%). Analysis of 36 *Trpm4-L903* ABE9-SpRY edited mice showed a similar trend (**Fig. 5d,e**). The product purity was comparable, as observed for *Tpc1-I486*, at 93.9 ± 10.3% [median (IQR), 97.6% (97.4-97.8%)] (**Fig. 5f**), due to the limited bystander editing events at A8 (**Fig. 5c**). Overall, p.L903P was accurately and precisely established at even higher efficiencies than p.I486T, at frequencies of 28.3 ± 20.8% [median (IQR), 21.2% (10.4-34.7%)], and up to 96.0% (**Fig.5e**). Similarly, indel frequencies were slightly higher at the *Trpm4-L903* locus at 1.0 ± 5.9%, reaching 35.5% in one instance.

**Figure 5.**
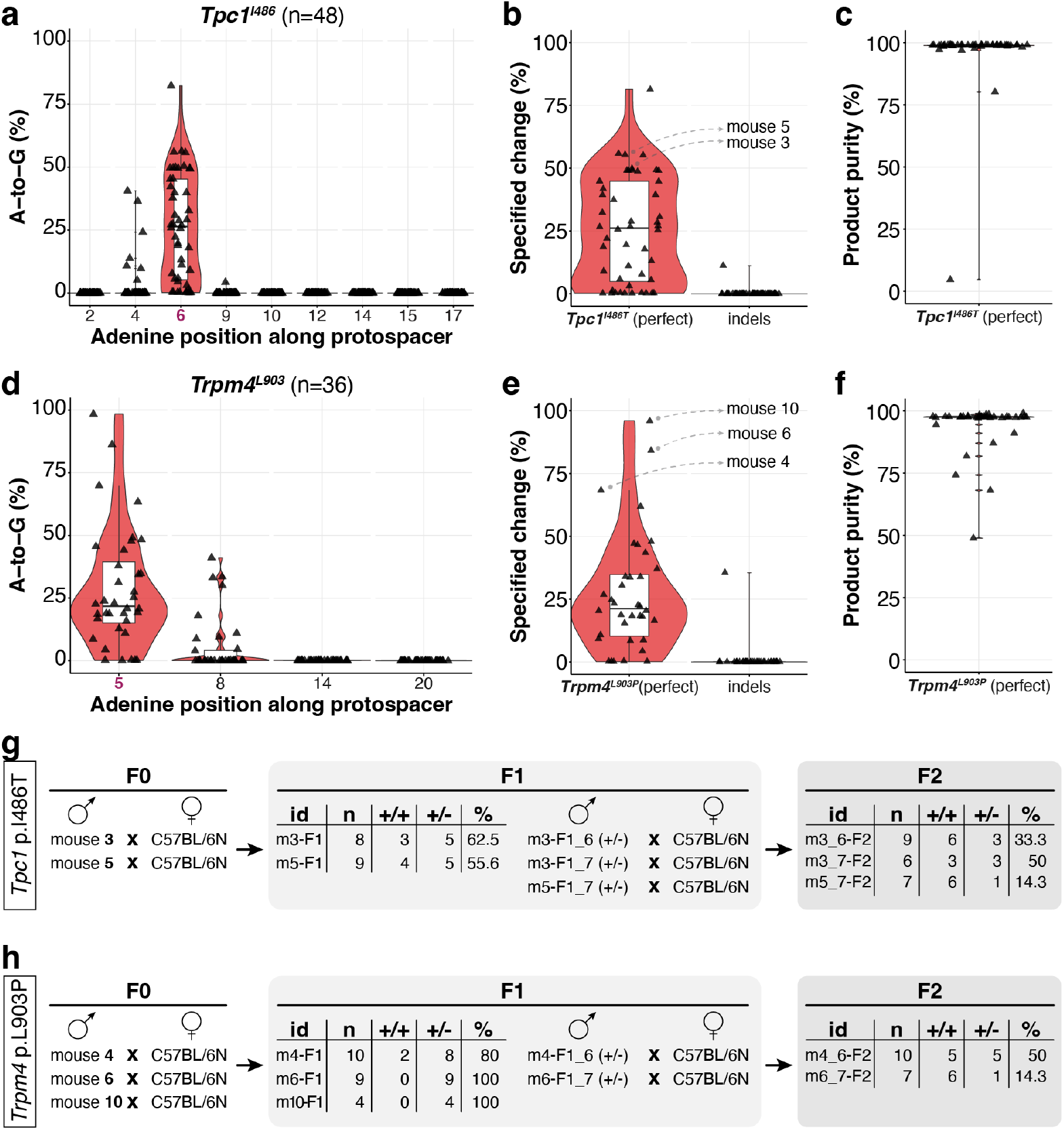
ABE9-SpRY can effectively produce *Tpc1*^*I486T*^ and *Trpm4*^*L903P*^ founder mice. A-to-G editing frequencies plotted across each adenine position are shown in (**a**) for 48 *Tpc1*^*I486*^ adult mouse biopsies, and (**d**) for 36 *Trpm4*^*L903*^ adult mouse biopsies. The frequency of desired editing (without bystander edits) and indels are presented in (**b**) for *Tpc1*^*I486*^ adults, and (**e**) for *Trpm4*^*L903*^ adult mice. Product purity is calculated as the percentage of correctly edited sequences relative to the total number of modified alleles and is displayed in (**c**) for *Tpc1*^*I486*^ adults, and (**f**) for *Trpm4*^*L903*^ adult mice. Data are presented as standard boxplots (white) following the Tukey convention, which includes the median (black horizontal line), displayed within violin plots (red), and individual data points (black) representing each biological replicate. Crossing schemes of F0 founder mice, with numerical data for F1 and F2 transmission for *Tpc1*^*I486T*^ (**g**), and *Trpm4*^*L903P*^ (**h**), respectively.

To determine the F1 transmission rate and establish stable mutant lines for *Tpc1* p.I486T and *Trpm4* p.L903P, we proceeded to mate two verified adult F0 *Tpc1* p.I486T male mice **(Fig. 5b)** and three verified adult F0 *Trpm4* p.L903P male mice **(Fig. 5e)** with wild-type females. We observed high transmission rates for *Tpc1* p.I486T of 55.6% and 62.5% in the two independent matings **(Fig. 5g, Supplementary Fig. 7a)**, and high to complete transmission for *Trpm4* p.L903P at rates ranging from 80% to 100% **(Fig. 5h, Supplementary Fig. 7b)**. Finally, we mated select heterozygous F1 carriers with wild-type mice to establish stable lines (**Supplementary Fig. 8)**.

These results suggest that the pooled embryo analysis has somewhat underestimated the potential editing efficiency of ABE9-SpRY when deployed to individual loci rather than diluted across several for screening. Moreover, the highly precise editing rates and encouraging off-target profile indicate the potential for direct founder-generation phenotyping, which could accelerate experimental timelines and reduce both resource consumption and the number of animals required.

### Precise hiPSC engineering with ABE9-SpRY

Over the past two decades, hiPSCs have become a powerful *in vitro* model, offering a human-specific system for studying disease mechanisms, drug responses, and genetic variants in a physiologically relevant context when paired with directed-differentiation procedures ^55,56^. Given this importance, we aimed to validate the potential of ABE9-SpRY for disease modelling in hiPSCs. To enrich base-edited cells, we used the XMAS-TREE assay ^57^, in which a successful A-to-G conversion of a stop codon in between an episomal mCherry-linker-EGFP construct leads to the co-expression of GFP alongside the pre-existing mCherry signal (**Fig. 6a**). We selected the human p.I485T mutation of human *TPC1*, corresponding to mouse *Tpc1* p.I486T, as a test and designed two sgRNAs with the target adenine located either in position A5 (sgRNA#2) or A6 (sgRNA#1) of the protospacer sequence, respectively (**Fig. 6b**). We then co-transfected either ABE8e-SpRY or ABE9-SpRY plasmid, which we re-cloned under an EF1α promoter for robust hiPSC expression, along with the reporter plasmid, a reporter-STOP-sgRNA plasmid, and either of the two target sgRNA plasmids, thereby enriching the base-edited cell population through FACS sorting (**Fig. 6c**). While the overall editing efficiency was substantially higher with ABE8e-SpRY, 75.8 ± 1.0% editing with sgRNA#1 and 33.7 ± 2.0% with sgRNA#2, ABE8e-SpRY was also active very broadly. In particular, the two adjacent bases A4 and A6, or A3 and A5 for sgRNA#1 and sgRNA#2, respectively, displayed very similar activity. By contrast, ABE9-SpRY achieved efficiencies of only up to 10.9 ± 0.7% (**Fig. 6d**). However, we observed superior overall desired editing activity without unwanted mutations, a result of the higher precision, with sgRNA#1 reaching 9.9 ± 0.7%. Notably, despite ideal target site positioning, sgRNA#2 showed a slightly lower efficiency at 7.6 ± 0.6% (**Fig. 6e**). The highest editing rate without unwanted mutations for ABE8e-SpRY was 3.0 ± 0.1%. While the overall editing rates of ABE9-SpRY were lower, this limitation is less critical in the context of hiPSC-based disease modelling, where correctly edited clones are typically isolated and expanded prior to downstream differentiation into specific cell types of interest. Importantly, such clonal isolation strategies rely on the high precision of the editing event, which we achieved with ABE9-SpRY, but not ABE8e-SpRY. In addition to the increased frequency of indels observed (**Fig. 6f**) at 2.9 ± 0.5% with ABE8e-SpRY compared to 0.1 ± 0.1% with ABE9-SpRY using sgRNA#1, these results further bolster the case for the potential of ABE9-SpRY in a wide variety of precision disease modelling applications, spanning from animal to cellular models.

**Figure 6.**
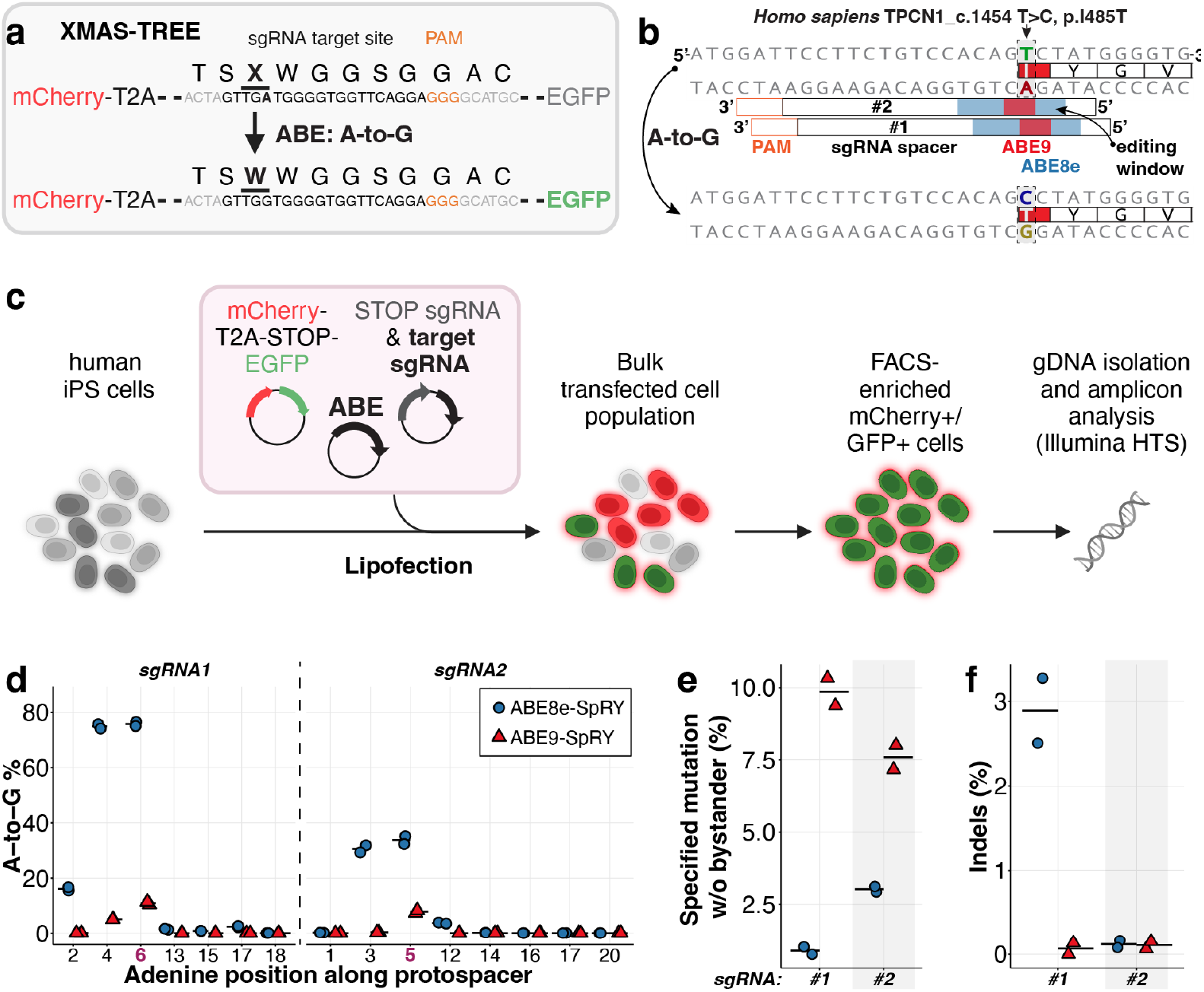
Enrichment and analysis of base-edited hiPSC populations via XMAS-TREE. (**a**) Diagram of the X-MAS TREE system, illustrating the expression of an mCherry cassette followed by a stop codon (TGA) and a GFP cassette. Conversion of the adenine in the stop codon to guanine activates GFP expression. (**b**) Diagram showing the tiling of two sgRNAs across the target genomic region. (**c**) Schematic of the X-MAS TREE workflow for identifying and enriching adenine base-edited cells. Flow cytometry was used to isolate populations that are double-positive for mCherry and GFP for downstream sequencing. (**d**) Percentage of adenine-to-guanine conversions along the protospacer; (**e**) percentage of desired edits without bystander effects; and (**f**) percentage of indels observed with ABE8e-SpRY or ABE9-SpRY, using sgRNA1 or sgRNA2 in sorted cells. Horizontal bars represent the mean from two biological replicates. Parts of the schematics in **c** were created with BioRender.com.

## DISCUSSION

Base editing holds great promise for modelling point mutations due to its benefits over existing techniques, as it prevents DSBs and shows high on-target efficiency ^12,13^. While previous studies have utilised base editing for animal modelling, they have frequently encountered issues such as off-target effects, bystander editing, and indels ^47,48,58,59^.

Successful disease modelling via base editing requires (1) precise editing with no bystander missense mutations, as they can easily mask or disrupt the interpretation of the desired mutation ^60^; (2) absence of indels on the same allele to avoid frameshifts and truncated proteins ^61^; and (3) minimal off-target effects, which, if frequent, can persist despite outcrossing ^62^ and may necessitate costly genome-wide validation.

Here, we utilised the latest advancements in the gene editing toolbox to address these issues. We demonstrated that using ABE9, an adenine base editor recognised for its high specificity and safety profile ^21^, fused with the PAM-less SpRY-Cas9 nickase ^22^, can efficiently introduce A-to-G mutations with great flexibility. It results in low to no bystander editing, minimal detectable Cas-dependent off-target effects at only two sites, and very few indels, achieving up to 89% precise editing of desired changes in mouse embryos. Moreover, in adult F0 founder mice, we observed up to 96% desired editing at the *Trpm4-L903* locus, highlighting the potential for phenotyping within the same generation. In contrast, ABE8e-SpRY exhibited a higher overall A-to-G conversion efficiency, close to homozygosity. However, this was accompanied by bystander mutations at similar frequencies, resulting in significantly lower product purities. Furthermore, ABE8e-SpRY caused substantial levels of Cas-dependent off-target editing in mouse embryos at 14 of 20 sites examined, with A-to-G editing frequencies at OT5 of the *Trpm4-L903* sgRNA reaching up to 84%. Therefore, the trade-off between reduced editing efficiency and high precision remains a valuable advantage that ABE9-SpRY offers for generating genetic and disease models where bystander editing is unacceptable.

We found that pooling sgRNAs during the initial *in vivo* testing in embryos is an effective strategy. This approach enables rapid screening of editing outcomes with reduced effort and decreases the number of animals required. However, this approach seems to underestimate editing efficiency compared to experiments where only a single sgRNA is injected. For example, at the *Tpc1-I486* locus, perfect editing (desired editing without bystander edits) occurred at a frequency of 10.7 ± 17.5% in embryos (pooled) versus 25.6 ± 21.0% in adults (single). Similarly, at the *Trpm4-L903* locus, the desired editing frequency was 21.3 ± 28.4% (pooled) versus 28.3 ± 20.8% (single). This discrepancy may stem from the dilution of the editing machinery across four different target sites, and potentially their off-targets, in the pooled condition. This likely reduces the effective concentration of the editor at each individual locus, thereby lowering the observed editing efficiency.

At the *Tpc2-K188* locus, ABE8e-SpRY demonstrated efficient editing, while only one mouse embryo showed any editing for ABE9-SpRY. This indicates that the SpRY-Cas9n protein can access this target site, but the evolved TadA-9 deaminase enzyme has shown reduced substrate acceptance compared to TadA-8e. Given that the N108Q and L145T mutations in ABE9 are speculated to expel the backbone of the DNA substrate ^21^, this may also change the substrate-interaction behaviour of TadA-9 compared to TadA-8e, in addition to altering the editing window. As such, the observation at the *Tpc2-K188* locus suggests that, in addition to altering the editing window, these mutations may create sequence-dependent steric hindrances that prevent the enzyme from effectively engaging with the target adenine, even within its supposedly optimal editing window. To better understand the genome-wide targeting properties of ABE9-SpRY, a combination of detailed structural analyses to elucidate the mechanisms by which the N108Q and L145T mutations influence substrate binding and high-throughput *in vitro* analysis of a large number of disparate target sites will be required. However, despite its efficiency, ABE8e-SpRY would not be the alternative to ABE9-SpRY in cases such as those at the *Tpc2-K188* locus due to its off-target profile. For sites untargetable by ABE9-SpRY, alternative variants with enhanced specificity, such as ABE8e-WA/WQ ^30^, ABE8e-N108Q ^21^, ABE8e-V106W ^20^, and next-generation ABEs (e.g., ABE8r, ABE-Umax, ABEx2/4, E2-/E4-ABE, TadA-8e-L1B_S116, hpABE5.20 ^63-68^, are emerging and may serve as effective substitutes when fused to SpRY-nickase. In practice, the advent of scalable machine learning-based engineering tools to create bespoke ABE-SpCas9 variants with target-specific PAMs ^69^ will help to maximise on-target editing by use of the highest activity and precise deaminases with context-specific PAM-interacting motifs, to substantially reduce off-target editing.

The PAM-less properties of SpRY, in any combination with next-generation genome editors, present significant challenges of their own, particularly concerning increased off-target genome editing, which consequently leads to a rise in chromosomal rearrangements ^11^. As a result, ABE-SpRY editors are expected to demonstrate increased off-target editing due to their ability to bind and modify DNA sequences without the stringent necessity for a PAM. ABE8e-SpRY demonstrated extensive Cas9-dependent off-target editing at 70% of the examined off-target sites. At one of these sites, *Trpm4-L903* OT5, editing reached up to 84%. ABE9-SpRY, in contrast, exhibited editing at only 10% of the examined sites. We acknowledge that a limitation of our off-target analysis is its dependence on *in silico* predictions of potential non-specific sgRNA binding sites. This method may introduce bias, as the sites are selected *a priori* and might miss *bona fide* off-target effects. Nonetheless, it remains a common and practical initial method for evaluating base editor specificity. To complement *in silico* predictions, a suite of unbiased genome-wide off-target detection methods has been developed in recent years ^70^. For adenine base editors, specific unbiased genome assays like EndoV-seq (76), Digenome-seq (77), and Selicit-seq (78) have been developed. Especially when stringent validation is necessary, such as in clinical settings, these should be complemented with detailed chromosome-level structural analyses using assays like CAST-seq (80). Although these advanced methods are valuable, our results using *in silico* prediction already demonstrate that ABE9-SpRY strikes an attractive balance of precision and flexibility, supporting its suitability for generating disease models.

hiPSCs are increasingly utilised for disease modelling because they closely resemble the human genome and can differentiate into specific tissues ^71^. As hiPS cell culture continues to advance from traditional 2D systems to more intricate 3D models, it enables advanced exploration of pathophysiological mechanisms and disease modelling ^72^. For this purpose, reproducing genetic variants as precisely as possible is crucial, making the exclusion of bystander editing and limitation of off-target effects indispensable ^73^. Given the target flexibility of ABE9-SpRY, we designed two tiled sgRNAs across the target site and demonstrated that ABE9-SpRY could precisely induce a single-base change in hiPSCs without bystander editing. Although editing efficiency is relatively low compared to ABE8e-SpRY, and clonal isolation remains necessary for certain downstream gene mutation studies, the absence of bystander editing simplifies the clonal isolation. Utilising reporter assays for editing enrichment, such as XMAS-TREE, and optimising the delivery method will help overcome the limitations of low editing efficiencies ^74,75^.

Overall, ABE9-SpRY demonstrates high precision while maintaining considerable flexibility, which is crucial, as missense mutations can be highly detrimental. For example, in the context of disease rescue, even with ABE8e’s high efficiency in introducing the intended point mutation, a bystander edit caused a missense mutation that ultimately prevented phenotypic rescue ^60^. This underscores the importance of precision editors like ABE9-SpRY, particularly when modelling diseases or designing therapeutic strategies where functional outcomes are paramount. This level of precision is essential for fundamental research, such as structure-function studies of a specific protein domain, where even a single unintended amino acid change introduced by bystander editing could make the results uninterpretable. The potential for base editors to introduce confounding SNVs, as systematically demonstrated in mouse embryos ^59^, underscores the need for cleaner tools like ABE9-SpRY to produce reliable models for functional analysis. With ABE9-SpRY’s versatility in targeting specific adenines, along with minimised bystander editing and reduced Cas-dependent and Cas-independent off-target effects, it represents a valuable tool for creating precise disease models in both animals and cell lines, hiPSCs, and organoids derived from them.

## METHODS

All methods were performed in accordance with the relevant guidelines and regulations.

### Reagents and biological resources

Information on major resources is provided in **Supplementary Table 1**.

### Mouse strains

Wild-type C57BL/6N mice were purchased from Charles River Laboratories (Wilmington, MA, USA). The experimental procedures were approved by the regional council of Karlsruhe, Germany, in accordance with the Animal Welfare Act (AZ35-9185.81/G-60/23). Microinjection was performed in the cytosol of C57BL/6N zygotes. All animals were housed in the Interfaculty Biomedical Faculty (IBF) of Heidelberg University. Mice were maintained under specified pathogen-free conditions with a 12-hour light/12-hour dark cycle, and water and standard food (Rod18, LASvendi GmbH, Germany) available ad libitum. All mouse experiments described in this study comply with ARRIVE guidelines.

### Identification and Alignment of Mouse Orthologues for Targeted Point Mutations

To identify the corresponding mutations in the mouse genome, to four target loci of interest with missense mutations *Tpc1 p*.*I485T, Tpc2 p*.*L265P, Tpc2 p*.*K204A*, and *Trpm4 p*.*L907P* we retrieved the *Mus musculus* and *Homo sapiens* gene sequences from Ensembl and performed protein sequence alignments. This allowed us to pinpoint the target amino acids and the corresponding adenines in the mouse coding sequence. Through this alignment, we identified the orthologous amino acid residues in the respective mouse genes and the potential missense mutations we can model by A-to-G transition mutation mediated through ABEs: *Tpc1 p*.*I486T, Tpc2 p*.*L249P, Tpc2 p*.*K188G*, and *Trpm4 p*.*L903P*. Notably, for *Tpc2 p*.*K188G*, a glycine substitution was created *in silico* instead of the alanine. We decided to proceed with the glycine substitution based on the similar chemical and functional properties of glycine and alanine.

### Design and cloning of sgRNAs

sgRNAs were designed manually to enable the precise installation of the defined missense mutations and validated using ACEofBASEs ^53^ with the following parameters: select BaseEditor “custom ABE” with Window start “3” and Window end “11” with PAM “NRN” for *Tpc1-I486, Tpc2-K188*, and *Trpm4-L903* target sites or with PAM “NYN” for the *Tpc2-L249* target site; for Species “Mouse (Mus musculus) Ensembl V 103”. The full list of sgRNA oligonucleotides to clone and the sgRNAs used in this study are shown in Supplementary Tables 2 and 3.

For *in vitro* applications of sgRNAs, the 20 bp *Sp*Cas9 sgRNA spacer sequences used in this study were designed with the 5’ and 3’ sequences (**Supplementary Table 2**), as previously described ^76^, with an additional 5’ “G” added in cases in which the spacer does not start with a G. In brief, the 20 bp target site (N20) sequences were ordered as oligonucleotides with the following sequences 5’-CACC(N20-21)GTTTT-3’, while the sequences complementary to the target site were designed and ordered as oligonucleotides as follows 5’-CTCTAAAAC(N20-21)-3’. sgRNAs were cloned into the BsaI-digested 2.18 kb pU6-pegRNA-GG-acceptor fragment (pU6-pegRNA-GG was a gift from David Liu, Addgene plasmid #132777, ^76^) by oligonucleotide annealing and Golden Gate assembly as previously described ^76^, including the standard *Sp*Cas9 sgRNA scaffold. *R-loop SaCas9 sgRNAs*. To generate the *Sa*Cas9 sgRNA plasmid, BPK1520, containing a pU6_BsmBIcassette-Sp-sgRNA was used as a backbone (BPK1520 was a gift from Keith Joung, Addgene plasmid #65777, ^39^). The *Sp*Cas9 sgRNA scaffold was digested using *BsmBI* and *HindIII*, and the digested fragment was isolated. The *Sa*Cas9 sgRNA scaffold sequence ^42^, designed with complementary overhangs (**Supplementary Table 2**), was ligated into the digested vector backbone. The designed sgRNA spacer sequences were then ordered as oligonucleotides with *BsmBI* overhangs (**Supplementary Table 2**), and the annealed oligos were cloned into the *BsmBI*-digested *Sa*Cas9 sgRNA plasmid via Golden Gate assembly. In brief, in a 10 µl reaction volume, 50 ng of *Sa*Cas9 sgRNA plasmid, 10 units of *BsmBI*-v2 (NEB), 20 units of T4 DNA ligase (NEB), 1 µL of 1 µM annealed spacer oligonucleotides, 1X T4 DNA ligase buffer were assembled under the following thermal cycling conditions: digestion at 42°C for 4 minutes, followed by ligation at 16°C for 3 minutes, repeated for a total of 11 cycles. The reaction was then incubated at 80°C for 10 minutes to inactivate *BsmBI. hiPSC sgRNAs*. Cloning of targeting sgRNA into the pDT-sgRNA-XMAS-1x plasmid was performed as previously described (pDT-sgRNA-XMAS-1x was a gift from Xiao Wang, Addgene plasmid #164413, was performed as previously described ^57^).

All sgRNAs for mouse experiments were ordered via Integrated DNA Technologies (IDT) as custom Alt-R CRISPR-Cas9 sgRNA (**Supplementary Table 3**).

### Cloning of adenine base editor plasmids

All plasmids constructed here were cloned via Gibson Assembly using the NEBuilder HiFi DNA Assembly (NEB) kit with PCR products amplified from designated template plasmids with specific primers (**Supplementary Table 4**) with Q5 Hot Start DNA polymerase (NEB).

To obtain pCMV_ABE9, we used the ABE9 editor sequence form ^21^, and recloned pCMV_ABE8e (a gift from David Liu, Addgene plasmid #138489, ^20^) by introducing the two defining mutations N108Q and L145T by PCR, altogether assembled via four fragments (**Supplementary Table 4**): pCMV_backbone, ABE9_fragment_1_TadA*_Nterm_N108Q, ABE9_fragment_2_TadA*_N108Q_L145T, ABE9_fragment_3_Cas9n-Nterm. To clone pCMV_ABE8e-SpRY, we assembled the following PCR products: pCMV_backbone, ABE8e_fragment1_TadA*, and ABE9-SpRY_fragment_4_SpRY-Cas9n (amplified from pCMV-T7-ABE8e-nSpRY-P2A-EGFP (KAC1069), a gift from Benjamin Kleinstiver, Addgene plasmid #185912, ^77^). Similarly, pCMV_ABE9-SpRY was constructed by assembling: pCMV_backbone, ABE9_fragment_1_TadA*_Nterm_N108Q, ABE9_fragment_2_TadA*_N108Q_L145T, ABE9_fragment_3_Cas9n-Nterm, and ABE9-SpRY_fragment_4_SpRY-Cas9n.

For the transfection of hiPSCs, pCMV_ABE8e-SpRY and pCMV_ABE9-SpRY were recloned to be expressed under the control of the EF1α promoter. In brief, two fragments were amplified, first the pEF1α_backbone (from pEF1α-hMLH1dn, a gift from David Liu, Addgene Plasmid #174824, ^78^) and then either ABE8e-SpRY or ABE9-SpRY, for subsequent Gibson assembly cloning.

The entire sequence of all constructs assembled using PCR was verified with the PlasmidEZ (Azenta/GeneWiz) full-plasmid sequencing service. All plasmids used for the transfection experiments were prepared using the EndoFree Plasmid Maxi Kit (Qiagen).

### Generation of adenine base editor mRNA

mRNA for the microinjection of mouse zygotes for ABE8e-SpRY and ABE9-SpRY was generated by *in vitro* transcription from *SalI*-HF linearised pCMV_ABE8e-SpRY and pCMV_ABE9-SpRY plasmid templates, respectively, using the HiScribe T7 ARCA mRNA Kit. The mRNA was purified using the RNeasy Mini Kit. mRNA quality was confirmed using the TapeStation RNA ScreenTape.

### Mammalian cell culture and transfection

Human HEK 293T cells and Neuro2a cells were purchased from ATCC and maintained in Dulbecco’s modified Eagle’s medium (DMEM; Gibco) supplemented with 10% FBS (Gibco) at 37 °C with 5 % CO_2_, up to 20 passages, by passaging cells every two to three days. hiPSCs were purchased from Greenstone Biosciences and cultured on hESC-Qualified Matrix Matrigel-coated (Corning) 6-well plates (Sarstedt) in StemMACS PSC-Brew XF medium (Miltenyi Biotec). Medium was changed daily, and cells were passaged every three days using StemMACS Passaging Solution XF (Miltenyi Biotec). The culture medium was supplemented with 2.5 µM Rock Inhibitor (Y27632, Miltenyi Biotec) on passaging days for 24 hours. The supernatant media from cell cultures were routinely analysed for mycoplasma presence using the MycoSPY Master Mix (Biontex).

All HEK293T and N2a transfection experiments were conducted with at least three independent biological replicates. HEK293T and N2a cells were seeded at a density of roughly 110,000 and 180,000 cells per well in 24-well plates, respectively, 20-24 hours before transfections. Transfections were performed with 750 ng of base editor plasmid, 250 ng of sgRNA expression plasmid, and, for experiments involving GFP co-transfection, 200 ng of pMAX-GFP (Lonza), combined with 2 μl of Lipofectamine 2000 (Thermo Fisher Scientific) in a total volume of 50 μl Opti-MEM (Thermo Fisher Scientific), according to the manufacturer’s instructions. Medium was exchanged to culture medium approximately 24 hours after transfection. For the Orthogonal R-loop assay, HEK293T cells were seeded at a density of 110,000 cells per well in 24-well plates, as described above. However, the cells were co-transfected with a mixture containing 500 ng of plasmids encoding base editor plasmids, 350 ng of *Sp*Cas9 sgRNA (*HEKsite2*), 500 ng of d*Sa*Cas9, 350 ng of *sa*Cas9 sgRNAs targetting separate genomic loci unrelated to the on-target site, and 200 ng of pMAX-GFP (Lonza).

hiPSCs were seeded on 6-well plates and transfected after approximately 24 hours at around 60 % confluency. Before transfections, the medium was changed to StemMACS PSC-Brew XF supplemented with StemMACS PSC-Support XF (Miltenyi Biotec). Transfections were performed with 300 ng pEF1α-XMAS-1xStop (a gift from Xiao Wang, Addgene plasmid #164411, ^57^), 300 ng pU6-SpCas9-XMAS-1x plasmid, 2100 ng of pEF1α_ABE8e-SpRY or pEF1α_ABE9-SpRY, and 10 µl Lipofectamine Stem (Thermo Fisher Scientific) per well, according to the manufacturer’s instructions. Medium was exchanged to culture medium approximately 24 hours after transfection.

### Fluorescence-activated cell sorting

Approximately 48 hours after transfections, HEK293T, N2a or hiPSCs were harvested for sorting. HEK293T and N2a cells were treated with TrypLE Express Enzyme (Thermo Fisher Scientific) to singularise the cells, followed by a washing step with D-PBS. The cells were resuspended in buffer containing 1x PBS without Mg^2+^/Ca^2+^ (Thermo Fisher Scientific), 0.5% FCS (Thermo Fisher Scientific) and 1% Penicillin-Streptomycin and filtered through a 35 µm cell strainer (Corning). hiPSCs were dissociated with Accutase (Thermo Fisher Scientific) and washed with D-PBS, followed by resuspending the cells in 1x PBS without Mg^2+^/Ca^2+^, 0.5% FCS, 2.5 µM Rock Inhibitor and passed through a 35 µm cell strainer (Corning). HEK293T and N2a cells co-transfected with pMAX-GFP were sorted for the top 30% GFP-positive population. For the XMAS-TREE reporter assay in hiPSCs, only cells co-expressing GFP and mCherry were sorted. FACS sorting was performed with either a BD FACSAriaIII or BD FACSAria Fusion at the Flow Cytometry & FACS Core Facility (FFCF), Zentrum für Molekulare Biologie der Universität Heidelberg (ZMBH) or FACS Core Facility (dFCCU), Uniklinikum Heidelberg, respectively.

### Genomic DNA extraction from mammalian cell culture

After FACS HEK293T, N2a cells were collected in 500 µl DMEM GlutaMAX supplemented with 10% FBS. hiPSCs were collected in 500 µl StemMACS PSC-Brew XF medium supplemented with 2.5 µM Rock Inhibitor. Cells were collected by centrifugation at 20,000 *x* g for 2 minutes, washed with 500 µl D-PBS, and lysed in mammalian gDNA lysis buffer (10 mM Tris-HCl, 0.05% SDS, pH 8, 800 units proteinase K (NEB)) for 2 hours at 37°C to extract genomic DNA. Afterwards, proteinase K was inactivated for 30 min at 80°C. Cells transfected without fluorophore plasmid were washed once 72 hours after transfection, and then lysed in mammalian gDNA lysis buffer, as described above.

### Microinjection of mouse zygotes

Injections were performed as previously described ^79^, with minor modifications. In brief, the injection mix contained adenine base editor mRNA (100 ng/μl) and sgRNAs (50 ng/μl each; 12.5 ng/µl each, for pooled sgRNA injections) in injection buffer (10 mM Tris-HCl, 0.25 mM EDTA, pH 7.4) to a final volume of 50 μl with molecular biology-grade water. To prevent clogging of the cannula, the injection mix was filtered shortly before microinjection using a Corning Costar Spin-X 0.22 μm centrifuge tube filter. Zygotes were collected from timed matings and cultured *in vitro* in M2 medium during the injection. Injection into the cytoplasm of zygote-stage embryos was conducted by carefully inserting the injection needle at the equatorial level into the cytoplasm of each embryo, delivering a volume of 1-2 picolitres. Subsequently, the embryos were placed in preincubated M16 medium to select lysed and intact embryos. Embryos were transferred on the same day via oviduct transfer to 0.5-day pseudopregnant foster mothers.

### Genomic DNA extraction from mouse samples

Embryonic day 14 mice were isolated from surrogate mothers. For DNA extraction, each embryo was sectioned into smaller pieces, and genomic DNA was isolated using the DNeasy Blood & Tissue Kit (Qiagen) following the manufacturer’s protocol, with minor modifications. All reagents were applied at double the recommended volume since the starting material exceeded 10 mg per embryo. Proteinase K from New England Biolabs (NEB) was utilised instead of the kit-supplied enzyme.

For genotyping of founder mice, ear biopsies were obtained from the Interfakultäre Biomedizinische Forschungseinrichtung of Heidelberg University. Genomic DNA was extracted from ear biopsy samples using a lysis buffer consisting of DirectPCR Lysis Reagent (Mouse Tail) (Viagen Biotech) and 0.25 µg/µl Proteinase K (AppliChem). For lysis, each sample was incubated in 100-150 µl of this lysis buffer at 55°C with vigorous shaking for a minimum of 4-5 hours, followed by inactivation of Proteinase K at 85°C for 45 minutes. The lysate is subsequently used directly for PCR.

### Off-target analysis in mouse embryos

The top five predicted off-target sites for each target locus were identified using ACEofBASEs ^53^ as described under sgRNA design (**Supplementary Table 7-10**). ACEofBASEs provides links to Ensembl gene IDs; therefore, the locus information for these sites was extracted from Ensembl. Importantly, the column “distance” in the off-target list was used when exporting Ensembl sequence information. In brief, when navigating the ENSEMBL genome browser, to export the locus information, ensure the value of distance and an additional 1kb is added to the “5’ Flanking sequence” and “3’ Flanking sequence”. Primers were designed (Primer3 2.3.7) to flank the predicted off-target loci.

### Targeted deep sequencing and data analysis

Samples were prepared for Amplicon-EZ-based NGS analysis (GeneWiz/Azenta Life Sciences) as previously described ^53^. All locus-specific primers contained the 5’ partial Illumina adapter sequences (Fwd: 5’-ACACTCTTTCCCTACACGACGCTCTTCCGATCT-3’, Rev: 5’-GACTGGAGTTCAGACGTGTGCTCTTCCGATCT-3’), and are listed in **Supplementary Table 5**. Typically, samples corresponding to different target loci from each biological replicate were pooled into a single sample. The pooled NGS data were demultiplexed by mapping to the respective reference sequences later.

For a higher level of sample multiplexing, a sequencing library with indexing primers that include unique i5 and i7 indexes, as well as P5 and P7 adapter sequences, was established. In brief, following the preparation of PCR amplicons with locus-specific primers as described above (PCR1), a second round of PCR was performed on these amplicons to barcode each sample, thereby allowing for the pooling of samples from both the same and different target sites (PCR2). The primer pairs for the second PCR contain complementary regions to the partial Illumina adapters along with the unique i5 and i7 index combinations (**Supplementary Table 6)**. For multiplexing, PCR1 samples were amplified using Q5 Hot Start DNA polymerase (NEB) for 27 cycles, validated by agarose gel electrophoresis of a subsample, and diluted between 1:5 and 1:20 based on gel band density. PCR2 was then set up at a volume of 20 µl, containing 1 µl of the diluted PCR1 product, 0.5 µM of each forward (i5) and reverse primer (i7) to create a combinatorial dual-index library, and Phusion U Green Multiplex PCR Master Mix (Thermo Fisher Scientific); it was amplified for an additional 13 cycles. Individual PCR2 amplification was validated by agarose gel electrophoresis of a subsample, and 24 PCR2 samples were pooled, run on a 1% agarose gel, with specific bands excised and cleaned up using the Monarch DNA Gel Extraction Kit (NEB), measured by Qubit 4 Fluorometer (Thermo) using the Qubit dsDNA BR Assay Kit, and pooled to equimolarity totalling 1 µg DNA. Samples were sequenced by GeneWiz (Azenta Life Sciences) using the Sequencing only service with a 5% PhiX spike-in on an Illumina MiSeq with 2 x 250 bp, paired-end sequencing, at a depth of approximately 10 million reads.

NGS data (in fastq.gz format) were processed and analysed using CRISPResso2 version 2.2.11 ^80^. In brief, two types of analyses were run in batch mode. First, all samples were analysed in base editing mode with the following custom input parameters: --min_average_read_quality, 30; -- quantification_window_size, 20; --quantification_window_center, -10; --base_editor_output. Downstream analysis was conducted using R version 4.2.1 in R Studio (packages: tidyverse, ggplot, ggpubr, cowplot), with data used to plot Adenines across the protospacer sourced from the “Nucleotide_percentage_summary_around_sgRNA.txt” output file. “Precision” (Supplementary Figure 1) was calculated as previously described ^21^ by dividing the second highest adenine-based editing frequency by the highest (A5 or A6); for this purpose, the indel frequency was extracted from the “CRISPRessoBatch_quantification_of_editing_frequency.txt” file and calculated as previously described ^53^.

Second, to quantify precise gene editing outcomes, the HDR mode was employed with the following custom input parameters: --min_average_read_quality, 30; --expected_hdr_amplicon_seq; --quantification_window_coordinates, “spacer-7nt (upstream)” – “spacer+PAM sequence (+3nt)”; --discard_indel_reads, TRUE. Downstream analysis was conducted using R, as described above, with data sourced from the “CRISPRessoBatch_quantification_of_editing_frequency.txt” file, with slight modifications following^81^. In short, for each sample’s HDR amplicon, values under “Reads_aligned_all_amplicons”, “Reads_aligned”, “Unmodified”, “Modified”, and “Discarded” (HDR) were collected; additionally, the values under “Discarded” (REF) from the Reference amplicon were obtained. Specific values were then calculated as follows:

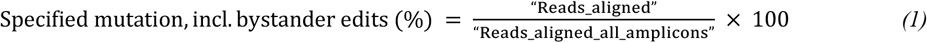

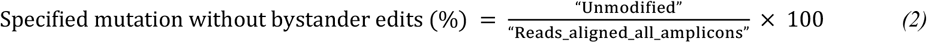

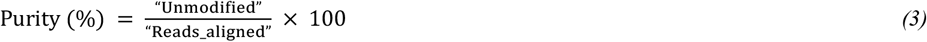

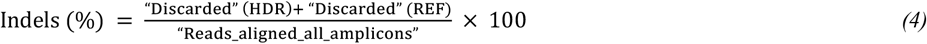

### Statistics and graphics

All data reported in the text are presented as mean ± standard deviation (SD). Where appropriate (n≥20), we also reported the median (Q2) and interquartile range (IQR) between the first quartile (Q1) and the third quartile (Q3). Statistical comparisons were conducted using two-tailed Welch’s *t*-test, as specified in the figure legends. Details on data reporting, including sample sizes and the statistical methods employed in each experiment, are provided in the figure legends. Data demonstrating statistical significance are depicted in the figures. Analysis and graphical data visualisation were performed in R version 4.2.1 with the Tidyverse, ggplot2, ggpubr, and cowplot packages ^82-85^. Figures were assembled in Adobe Illustrator, with some icons and schemes imported from Biorender.

## ACKNOWLEDGEMENTS

Sayari Bhunia is a member of HBIGS, the Heidelberg Biosciences International Graduate School. We thank the entire Freichel lab for their constructive feedback on the manuscript. We thank Maren Schneider for help with cloning pEF1α_ABE9-SpRY and XMAS-TREE sgRNAs. We thank the whole team from the Interfakultäre Biomedizinische Forschungseinrichtung (IBF) at Heidelberg University for their expert technical assistance. We thank Angela Wirth for writing the animal proposal. While preparing this work, the authors used AI-assisted technologies (Grammarly and ChatGPT) to improve readability and language.

## AUTHOR CONTRIBUTIONS

Jun Kai Ong: Conceptualisation, Data curation, Formal analysis, Investigation, Methodology, Validation, Visualisation, Writing – original draft, Writing – review & editing. Sayari Bhunia: Investigation, Validation, Writing – review & editing. Beate Hilbert: Investigation. Vanessa Kirschner: Investigation, Writing – review & editing. Sascha Duglosz: Investigation. Frank Zimmermann: Investigation. Marc Freichel: Conceptualisation, Funding acquisition, Resources, Writing – review & editing. Alex Cornean: Conceptualisation, Data curation, Formal analysis, Methodology, Project administration, Supervision, Visualisation, Writing – original draft, Writing – review & editing.

## SUPPLEMENTARY DATA

Supplementary Data will be made available online.

## CONFLICT OF INTEREST

Alex Cornean is a co-founder and shareholder of DataHarmony Ltd, a company that provides gene editing-related services and software solutions.

## FUNDING

This research was funded by the German Research foundation (DFG) through the Collaborative Research Centres CRC1550 (FKZ 464424253, projects B09 and S01), CRC1328 (335447717, project A21), DFG project FR 1638/5-1 (531047917), the DZHK (German Centre for Cardiovascular Research) and the BMBF (German Ministry of Education and Research); and The Health + Life Science Alliance Heidelberg Mannheim.

## DATA AVAILABILITY

Oligonucleotide and sgRNA sequences used in this study are listed in Supplementary Tables 2, 3, 5, and 6. All raw NGS sequencing data generated in this study were in the NCBI Sequence Read Archive (SRA) database as a BioProject with the accession number PRJNA1291991. pCMV_ABE9-SpRY and pEF1α_ABE9-SpRY will be made available through Addgene. Any other data and reagents will be made available upon reasonable request.

## Notes

### Competing Interest Statement

A.C. is a co-founder and shareholder of DataHarmony Ltd, a company that provides gene editing-related services and software solutions.

### Summary of Updates

Added F1 and F2 transmission data for Tpc1 and Trpm4 base editing (Figure 5 revised).

