## Supplementary Figures for "Precise generation of bystander-free mouse models with ABE9-SpRY"

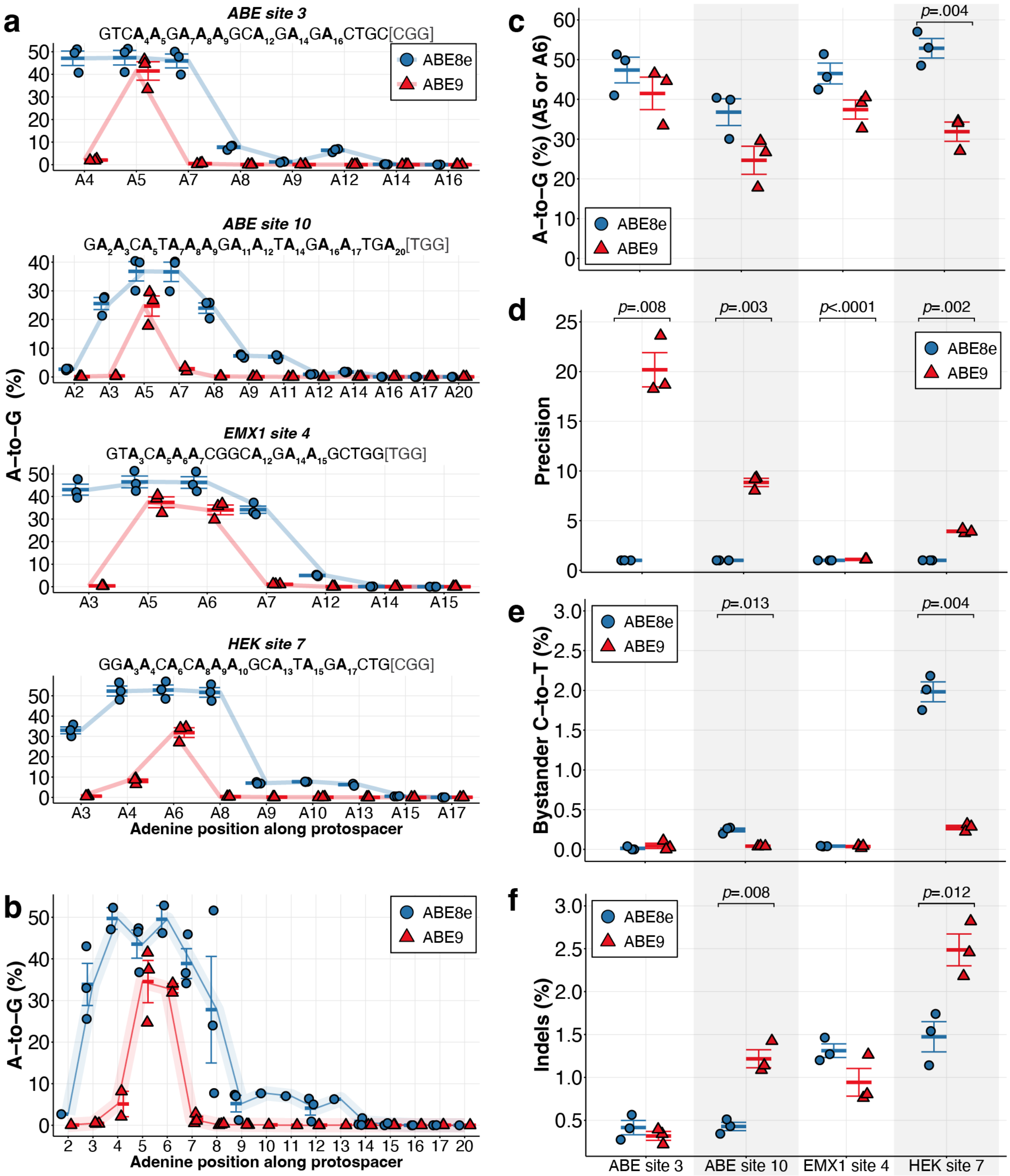


**Supplementary Figure 1 | ABE9's precision outweighs the higher A-to-G efficiency of ABE8e in human cells.** (**a**) Editing efficiencies of ABE8e and ABE9 across four endogenous target sites. (**b**) Summary plot of editing window across the four loci. (**c**) A-to-G editing frequency at adenine positions A5 or A6 (the highest is considered) at the four target sites. (**d**) Precision (according to Chen et al. 2023, *Nat Chem Biol*) calculated by dividing the highest A-to-G editing frequency at A5 (*ABE site 3, ABE site 10, EMX1 site 4)* or A6 (*HEK site 7*) by the second highest frequency at A4, A7, A6, and A4 for *ABE site 3, ABE site 10, EMX1 site 4*, and *HEK site 7*, respectively. (**e**) Unwanted C-to-T editing frequency measured across the protospacer sequence. (**f**) Indel frequency across the target sites. Error bars represent mean ± s.e.m. and individual data points for three independent biological replicates are shown. Statistical significance was determined by two-tailed Welch's *t*-test.

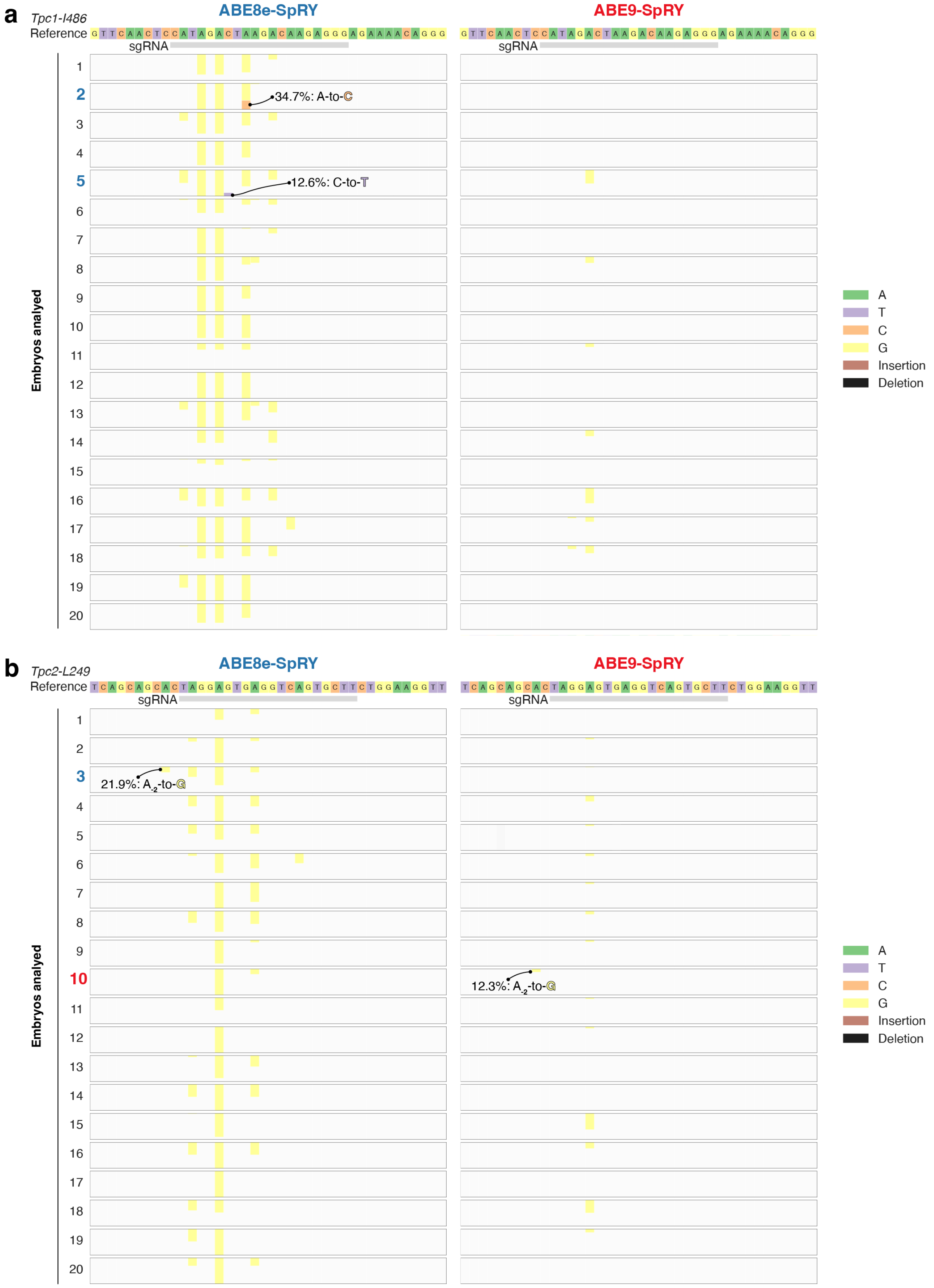


**Supplementary Figure 2 | Undesired on-target editing with ABE8e-SpRY and ABE9-SpRY at the *Tpc1$^{I486}$* (a) and *Tpc2$^{L249}$* (b) loci.** Graphs show the base composition in Crispresso2 plots of NGS analysis, highlighting, in colour, any modifications in edited embryos compared to the reference sequence 10 bp downstream and upstream of the sgRNA target site. The vertical size corresponds to the frequency of the modification.

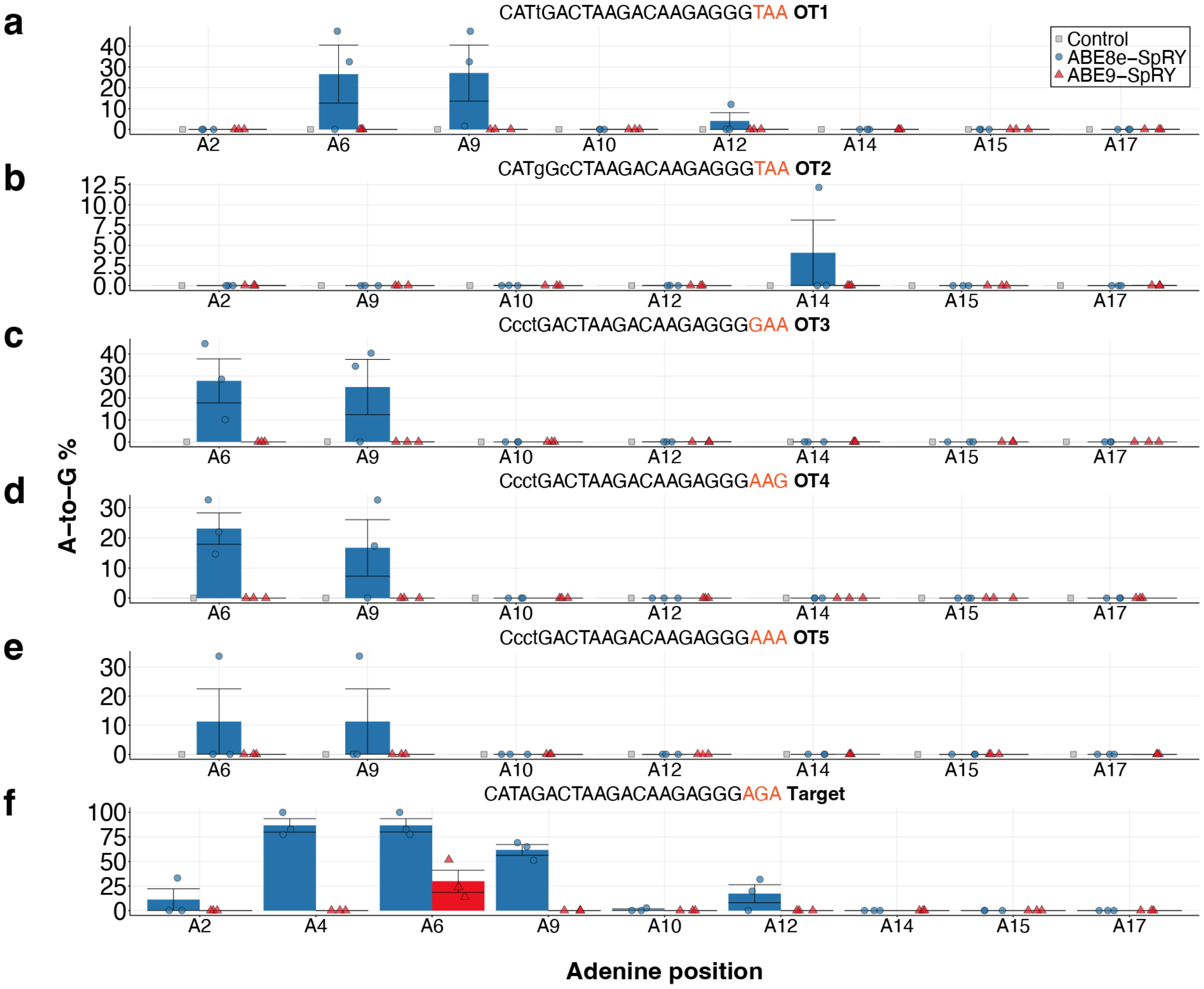


**Supplementary Figure 3** | ***Tpc1[I486]* off-target analysis.** (**a-e**) A-to-G frequency analysis of the top five predicted off-target site across all adenines within the protospacer sequence. Mismatches compared to the on-target sequence (**f**) are shown by lowercase letters while the PAM sequence is highlighted in orange. (**f**) Replotting of the on-target A-to-G editing efficiency for the embryos used for off-target evaluation (Fig. 3c). Error bars represent mean ± s.e.m. and individual data points for three independent biological replicates are shown.

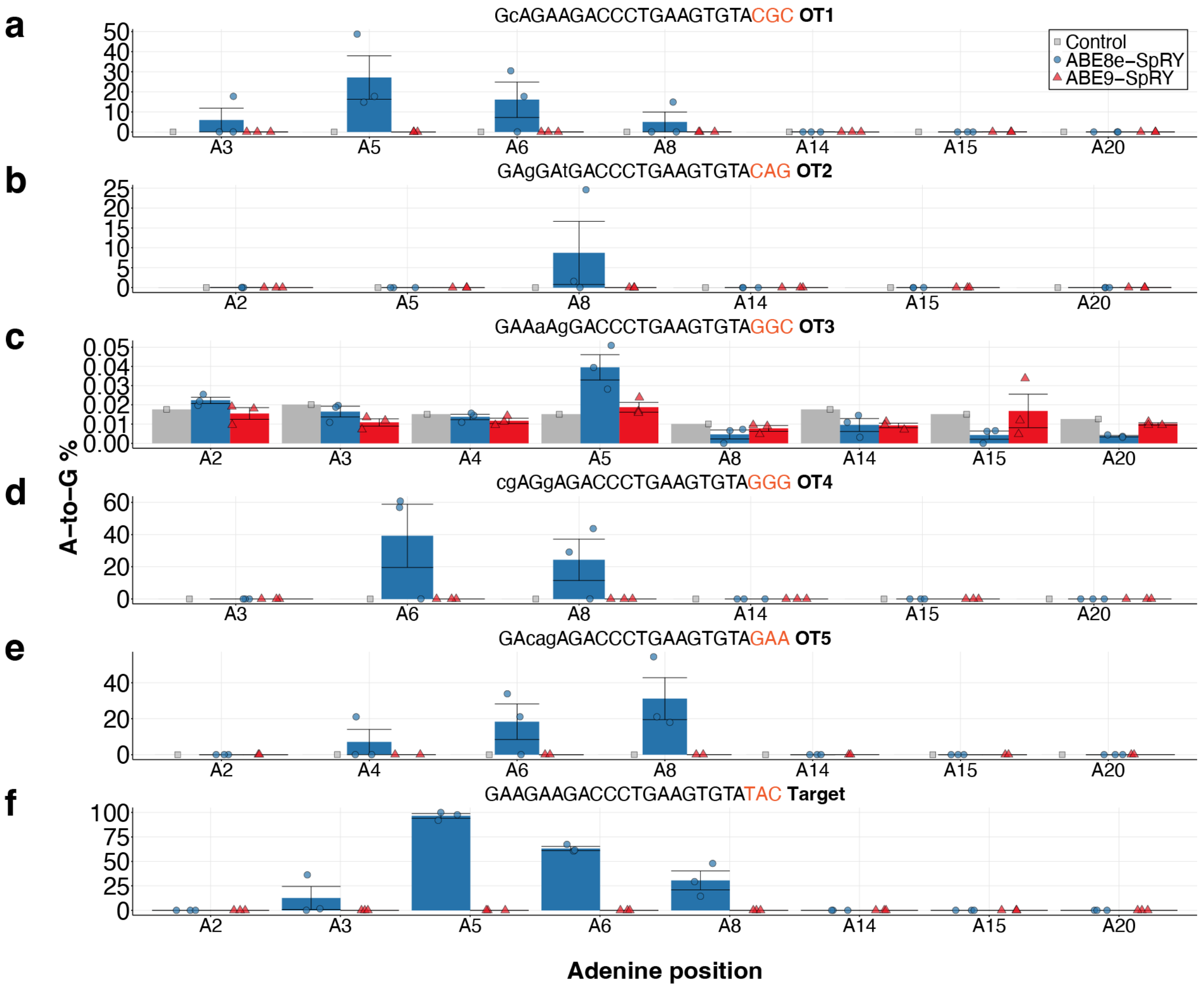


**Supplementary Figure 4 | *Tpc2*$^{K188}$ off-target analysis.** (**a-e**) A-to-G frequency analysis of the top five predicted off-target site across all adenines within the protospacer sequence. Mismatches compared to the on-target sequence (**f**) are shown by lowercase letters while the PAM sequence is highlighted in orange. (**f**) Replotting of the on-target A-to-G editing efficiency for the embryos used for off-target evaluation (Fig. 3c). Error bars represent mean ± s.e.m. and individual data points for three independent biological replicates are shown.

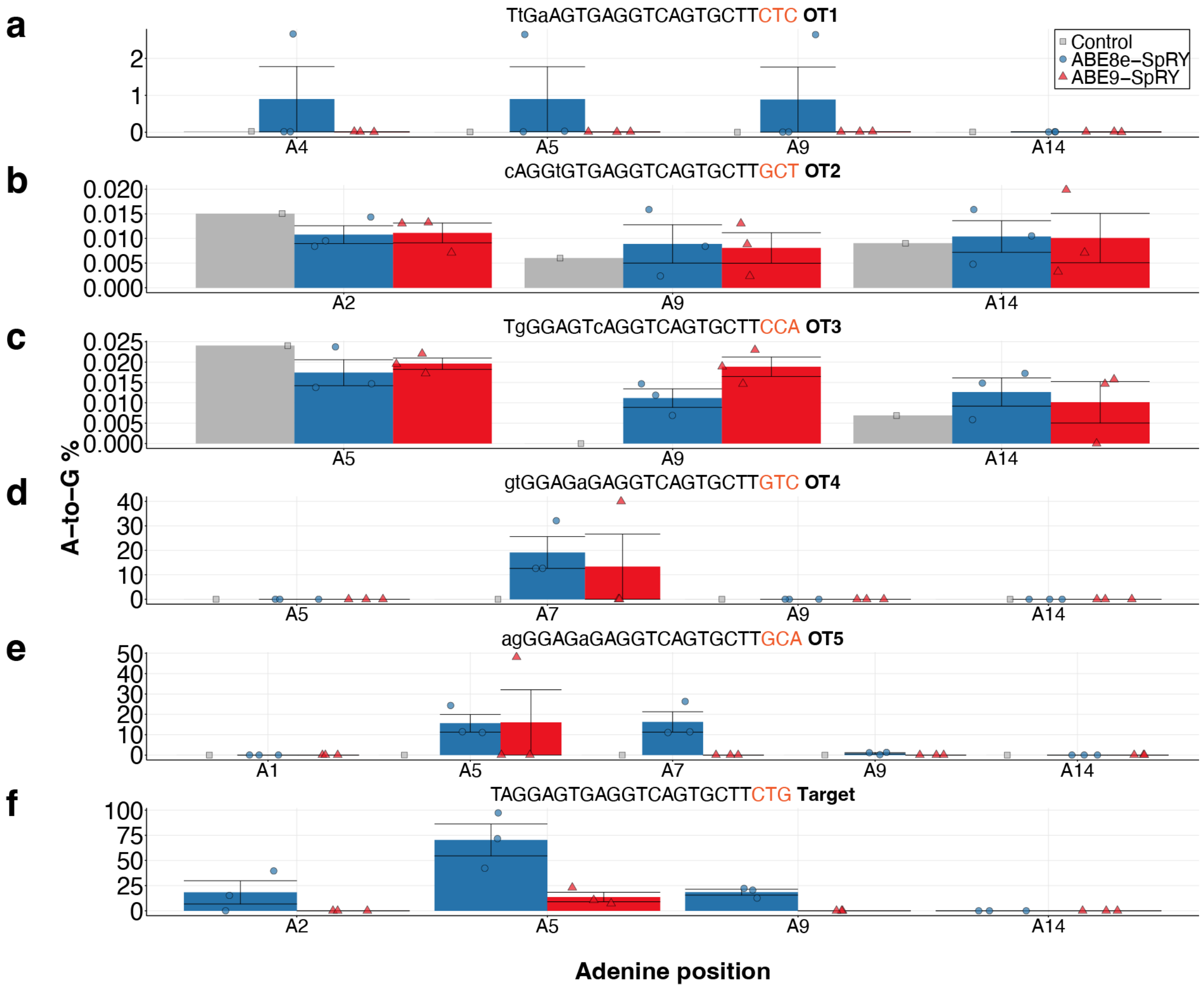


**Supplementary Figure 5 | *Tpc2$^{L249}$* off-target analysis.** (**a-e**) A-to-G frequency analysis of the top five predicted off-target site across all adenines within the protospacer sequence. Mismatches compared to the on-target sequence (**f**) are shown by lowercase letters while the PAM sequence is highlighted in orange. (**f**) Replotting of the on-target A-to-G editing efficiency for the embryos used for off-target evaluation (Fig. 3c). Error bars represent mean ± s.e.m. and individual data points for three independent biological replicates are shown.

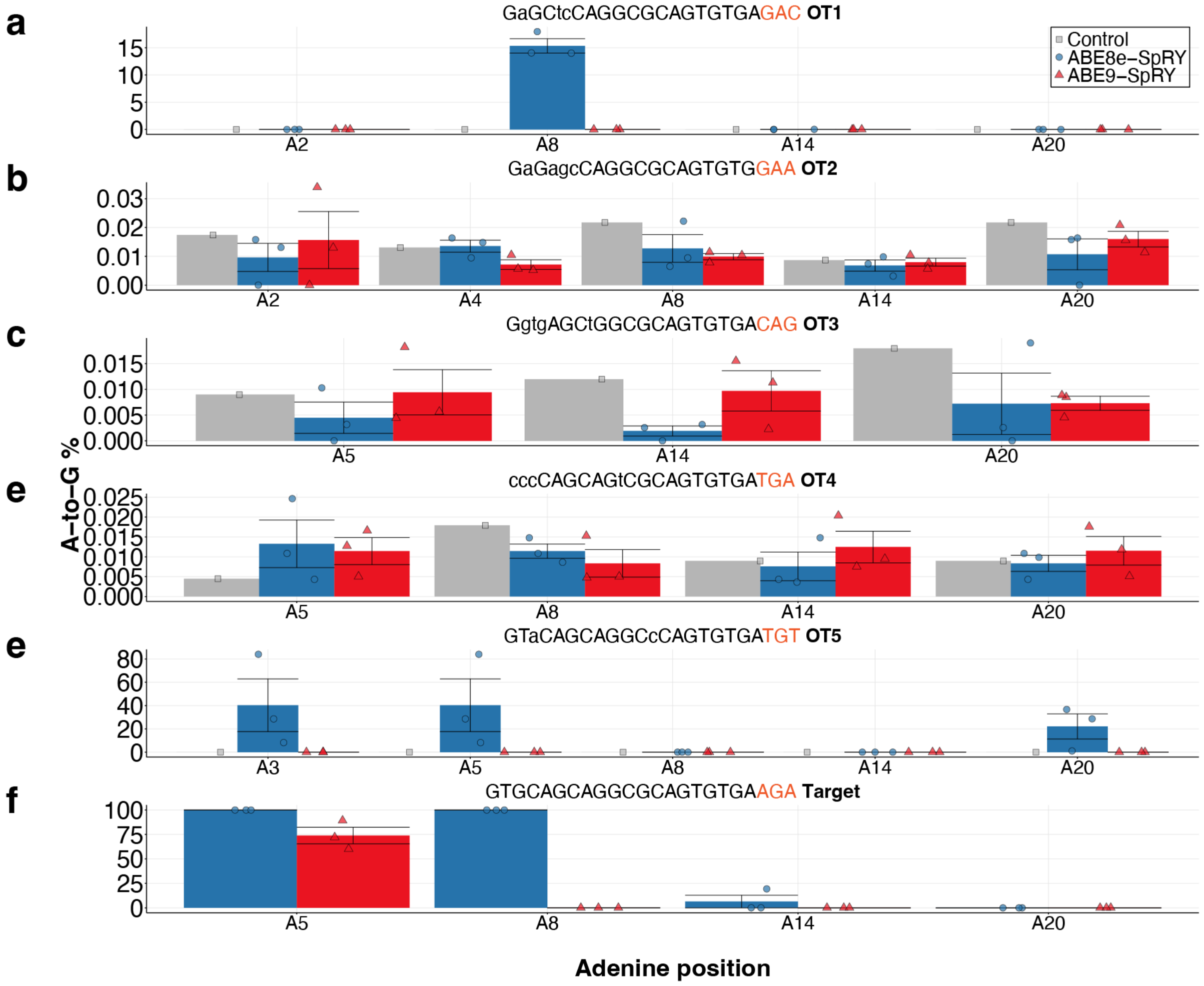


**Supplementary Figure 6 | *Trpm4*$^{L903}$ off-target analysis.** (**a-e**) A-to-G frequency analysis of the top five predicted off-target site across all adenines within the protospacer sequence. Mismatches compared to the on-target sequence (**f**) are shown by lowercase letters while the PAM sequence is highlighted in orange. (**f**) Replotting of the on-target A-to-G editing efficiency for the embryos used for off-target evaluation (Fig. 3c). Error bars represent mean ± s.e.m. and individual data points for three independent biological replicates are shown.

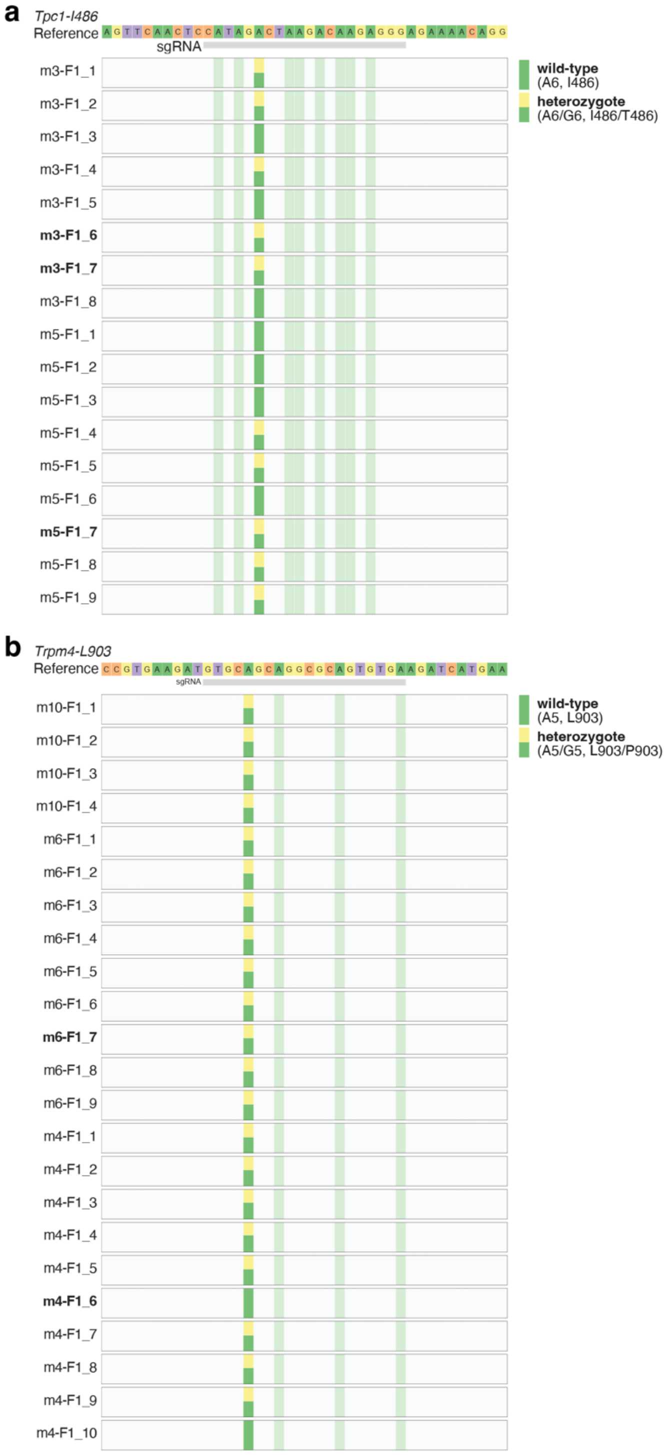


**Supplementary Figure 7 | Analysis of F1 transmission from ABE9-SpRY edited sites at the *Tpc1*$^{I486}$ (a) and *Trpm4*$^{L903}$ (b) loci.** Graphs show the base composition in Crispresso2 plots of NGS analysis, highlighting, in colour, any modifications in edited adult mice from ear biopsies compared to the reference sequence 10 bp downstream and upstream of the sgRNA target site. The vertical size corresponds to the frequency of the modification. A total for 17 (**a**) and 21 (**b**), adult mice were analysed to identify heterozygous carriers of the *Tpc1*$^{I486T}$ and *Trpm4*$^{L903P}$, respectively.

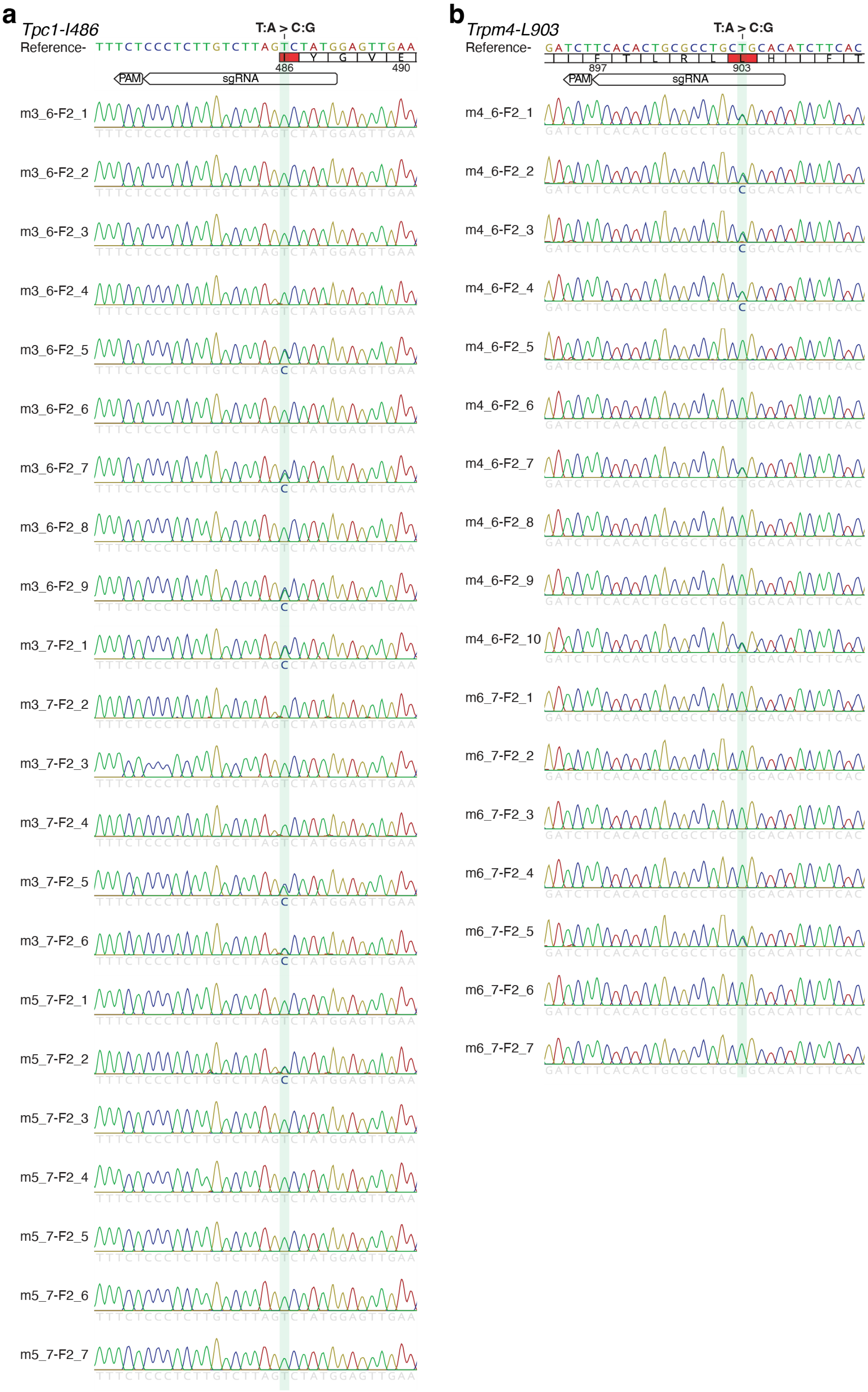


**Supplementary Figure 8 | Analysis of F2 transmission from ABE9-SpRY edited sites at the *Tpc1*$^{I486}$ (a) and *Tpc2*$^{L249}$ (b) loci.** Sanger sequencing reads for 22 and 17 F2 mouse ear biopsies were analysed for heterozygous carrier status of the *Tpc1*$^{I486T}$ and *Trpm4*$^{L903P}$ mutations, respectively. Note: for both targets, the sgRNA targets the complementary strand. Green shade highlights the mutated base.
